# Responders vs. non-responders to mesenchymal stromal cells in knee osteoarthritis patients: mechanistic correlates of donor cell attributes and putative patient features

**DOI:** 10.1101/2025.07.17.665028

**Authors:** Kevin P. Robb, Razieh Rabani, Shabana Vohra, Shoba Singh, Oreoluwa Kolade, Jaskarndip Chahal, Julie Audet, Anthony V. Perruccio, Ali Naraghi, Rajiv Gandhi, Osvaldo Espin-Garcia, Sowmya Viswanathan

## Abstract

Mesenchymal stromal cell (MSC) injection has afforded heterogenous outcomes in knee osteoarthritis (KOA). Herein, a framework that dually correlates KOA patient responsiveness to baseline autologous bone <u>m</u>arrow-derived MSC(M) donor batch attributes and baseline clinical and biomarker features is provided. Using clinical trial data, we demonstrated that MSC(M) with increased immunomodulatory potency are more efficacious. Multivariable MSC(M) genes correlated strongly with responder status and to 12- and 24-month improvements in Knee Injury and Osteoarthritis Outcome Scores. Responder MSC(M) donor batches had unique microRNA expression and ability to polarize CD14^+^ monocytes *in vitro*. KOA Responders had lower baseline physical activity and trended toward more severe baseline KOA. Baseline local but not systemic biomarkers showed trending correlations to patient responsiveness. 42% of KOA patients were Responders at 24 months, emphasizing durability of single MSC(M) injections. Together, our analytical methodology defines critical quality attributes of potent MSC(M) donor batches and identifies putative KOA patient theratypes to MSC treatments.

**Graphical Abstract:** 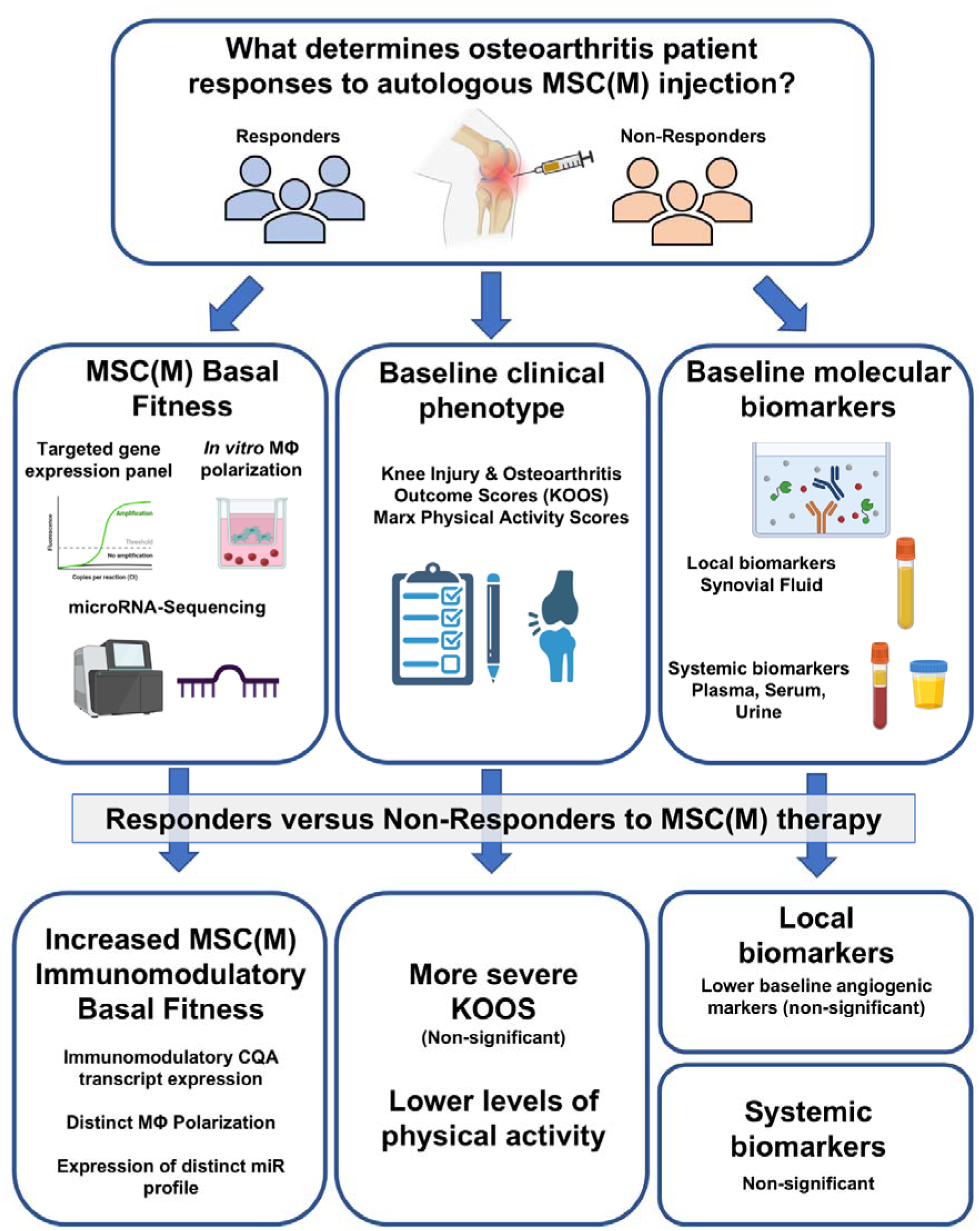

## INTRODUCTION

Knee osteoarthritis (KOA) affects 16% of adults over 40 years of age worldwide [1], with no available disease-modifying drugs. Pathophysiological mechanisms of KOA are heterogeneous and fluctuate temporally [2]. Culture-expanded mesenchymal stromal cells (MSCs) are being investigated for their multimodal mechanisms of action, including immunomodulatory functions [3,4]; however, there has been conflicting clinical efficacy reported. Equivocal results show no difference in 12-month improvements between umbilical cord-derived MSCs vs. first-line corticosteroids treatments [5]; conversely, other reports [6–9], meta-analyses [10,11], and randomized controlled trials [12,13] showed symptom and structural improvements with MSC treatments. This suggests heterogeneous responsiveness, stressing the need to identify signatures that distinguish responder from non-responder donor batches to deliver effective MSC treatments to KOA patients.

Heterogeneous responsiveness to MSC treatments is due to an interplay of both basal potency attributes of donor MSC batches and basal patient disease status [4]. Heterogeneity exists among MSC investigational products due to intrinsic variations in donors [4,14] and manufacturing parameters [15,16]; well-defined critical quality attributes (CQAs) that are sensitive, quantitative and relevant to disease-specific mechanism of action are thus needed [17].

Using reported [8], and new follow-up data from a phase I/IIa dose-escalation clinical trial, we conduct in-depth analyses of autologous donor bone <u>m</u>arrow-derived MSC (MSC(<u>M</u>)) batches and baseline/follow-up KOA patient characterization to provide an integrated algorithm to analyze heterogenous KOA patient responsiveness to MSC(M) treatments. We measured expression of a previously reported gene panel that correlated to *in vitro* functional activity [18], and determined that the gene panel significantly discriminated between responder vs. non-responders MSC(M) donor batches. MicroRNAs, important mediators of MSC function [19] further discriminated MSC(M) from Responders vs. Non-Responders.

KOA is recognized not as a single disease but a constellation of clinical phenotypes [20,21] and divergent molecular endotypes [22]. Here we report an exploratory analytical framework that combines clinical phenotypes and molecular endotypes that can be used as the basis for identifying KOA patient responsiveness to MSC(M) treatments.

## RESULTS

### Patient responder status

We systematically investigated multivariable factors affecting KOA patient responder status using 12- and 24-month patient-reported inverted KOOS data after injection with autologous MSC(M) [8]. We applied rigorous OMERACT-OARSI criteria [23] to determine responder status. Patients were classified as Function-Pain Responders vs. Non-Responders with Function-Pain Responders exceeding thresholds in <u>both</u> KOOS ADL and Pain at the 12-month time point. Non-Responders did not exceed thresholds in either ADL or Pain. Of twelve patients, six total Function-Pain Responders and six Non-Responders met these criteria at 12 months. While the sample size is limited, the binary criteria were useful in determining factors affecting patient clinical responses. However, correlations with all KOOS subscale scores were included to mitigate the limited power of binary analysis. There was no statistically significant dose-dependency [8]; Function-Pain Responders existed across all three MSC(M) doses including the 1 x 10^6^ (2/4), 10 x 10^6^ (1/4), and 50 x 10^6^ (3/4) cell doses, although more Function-Pain Responders were observed in the higher dose group (**Table 1**). We checked for associations between covariates (including sex, age, BMI, KL-grade, MSC dose and MRI baseline WORMs and synovitis scores) and Function-Pain Responder status and observed no significant correlations (**Table 1**).

**Table 1.** Summary of baseline patient characteristics. Association between each covariate were evaluated against responder status (p-values adjusted for multiple comparisons); no significant associations were detected. BMI: body mass index; KL: Kellgren-Lawrence; WORMS: Whole-Organ Magnetic Resonance Imaging Score [61].

| Covariate | Full Sample<br>(n=12 total) | Function-Pain<br>Responder<br>(n=6 total) | Non-Responder<br>(n=6 total) | p value | Adjusted<br>p-value |
| --- | --- | --- | --- | --- | --- |
| **Cell dose** |  |  |  | 0.7662 | 1 |
| 1x10<sup>6</sup> [n, (%)] | 4 (33) | 2 (33) | 2 (33) |  |  |
| 10 x10<sup>6</sup> [n, (%)] | 4 (33) | 1 (17) | 3 (50) |  |  |
| 50 x10<sup>6</sup> [n, (%)] | 4 (33) | 3 (50) | 1 (17) |  |  |
| **Gender** |  |  |  | 1 | 1 |
| Man x10<sup>6</sup> [n, (%)] | 7 (58) | 3 (50) | 4 (67) |  |  |
| Woman x10<sup>6</sup> [n, (%)] | 5 (42) | 3 (50) | 2 (33) |  |  |
| **Age** |  |  |  | 0.2289 | 1 |
| Mean (SD) | 54.8 (6.9) | 57.2 (4.5) | 52.3 (8.3) |  |  |
| Median (Min,Max) | 53.5 (45.0,65.0) | 56 (53,63) | 49 (45,65) |  |  |
| **BMI** |  |  |  | 0.5724 | 1 |
| Mean (SD) | 25.8 (3.5) | 25.4 (2.3) | 26.0 (4.6) |  |  |
| Median (Min,Max) | 25.5 (19.0,29.9) | 25.1 (23.3,29.7) | 27.9 (19.0,29.9) |  |  |
| **KL Grade** |  |  |  | 1 | 1 |
| 3 | 11 (92) | 6 (100) | 5 (83) |  |  |
| 4 | 1 (8) | 0 (0) | 1 (17) |  |  |
| **MRI Total Synovitis Score [65]** |  |  |  | 0.5199 | 1 |
| Mean (SD) | 6.8 (2.9) | 6.2 (1.9) | 7.8 (4.0) |  |  |
| Median (Min,Max) | 6 (3,12) | 5.5 (4.0,9.0) | 8 (3,12) |  |  |
| Missing | 2 | 0 | 2 |  |  |
| **MRI WORMS** |  |  |  | 0.3939 | 1 |
| Mean (SD) | 37.4 (9.3) | 34.8 (8.0) | 39.9 (10.4) |  |  |
| Median (Min,Max) | 36.2 (20.0,54.0) | 35.8 (20.0,44.0) | 41 (28,54) |  |  |

### 12-month Function-Pain Responder vs. Non-Responder classification holds at 24-months

Function-Pain Responders (categorized based on 12-month KOOS ADL and Pain) displayed significantly improved inverted KOOS scores in all subscales at 12- and 24-months follow-up (**Fig. 1A, Fig. S1**), supportive of persistent effects of a single injection of autologous MSC(M). Four of six Function-Pain Responders and five of six Non-Responders maintained their categorization between 12- and 24-months, indicating relatively stable and persistent clinical phenotypic changes (**Fig. 1B**). Two of six Function-Pain Responder and one of six Non-Responders changed their categorization based on changes in self-reported outcomes that are known to fluctuate [24]. Thus, at 24-months, five of twelve patients met Responder criteria while seven of twelve patients were Non-Responders. We report on Responder status based on 12-month data hereafter in this study.

**Fig. 1.**
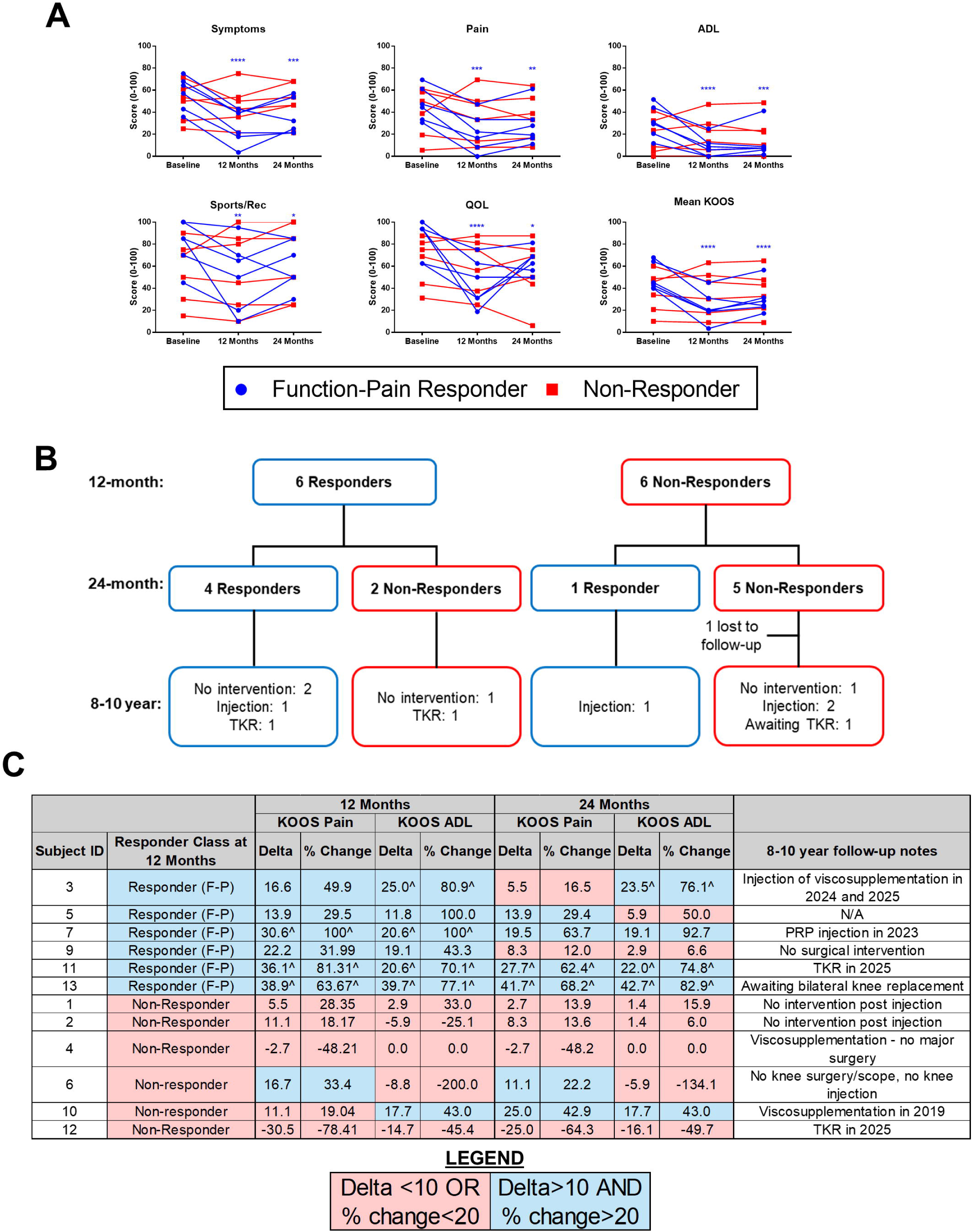
Osteoarthritis patient responders demonstrate improvements in KOOS subscales that are maintained out to 24 months follow-up post-injection. A) Patients categorized as Function-Pain Responders displayed significant improvements (indicated by lower score values) in KOOS subscales and mean KOOS at 12- and 24-months follow-up relative to baseline, while Non-Responders displayed no significant changes. KOOS scales were inverted such that higher scores represent greater knee-related issues. Two-way repeated measures ANOVA, Dunnett’s multiple comparisons test. *p<0.05, **p<0.01, ***p<0.001, ****p<0.0001; asterisks indicate significant differences for responder group relative to baseline at the time points indicated. B) Flow chart summary of patient responder status at 12-months, 24-months, and 8-10-years follow-up. C) Summary of OARSI-OMERACT Responder classification at 12 and 24 months, with 8-10 year follow-up notes. ^Indicates that patient exceeded substantial OARSI-OMERACT threshold of improvement in pain or in function ≥50% and absolute change (delta) ≥ 20. N=12 patients. F-P: Function-Pain Responder; ADL: function in daily living; Sports/Rec: function in sport and recreation; QOL: knee-related quality of life; KOOS: knee injury and osteoarthritis outcome score; PRP: platelet-rich plasma; TKR: total knee replacement.

We further collected patient follow-up at 8-10 years after a single MSC(M) injection. Of the six 12-month Function-Pain Responders, three received no interventions; one received viscosupplementation or platelet-rich plasma injections and two underwent knee arthroplasty. Of the six 12-month Non-Responders, one received no interventions; three received injections and one underwent knee arthroplasty (**Fig 1B,C**).

### Immunomodulatory gene panel discriminates between clinically effective and ineffective MSC(M) donor batches

Using biobanked autologous MSC(M) samples cultured under pro-inflammatory licensing conditions, we measured expression of a gene panel that correlated with *in vitro* immunomodulatory functionality [18]. Pro-inflammatory cytokine licensing conditions were used as a common method to measure immunomodulatory genes in MSC(M) [25].

Principal Component (PC) analysis revealed distinct unsupervised clustering between Function-Pain Responder vs. Non-Responder MSC(M) donor batches along the PC1 axis (29.3% of total variance) (**Fig. 2A**, **Fig. S2A**). Higher expression of *CCN2* and lower expression of angiogenic markers *THBS1*, *CXCL8*, and *ANGPT1* contributed most strongly to significantly lower PC1 scores observed in Function-Pain Responders (**Fig. 2B, Fig. S2A**). To ensure that individual donor batches were not skewing results, outlier analysis was performed by calculating T^2^ statistic, identifying three outlier data points (two Responder, one Non-Responder). However, the PCA with these data points removed, returned similar results; clear separation of Function-Pain Responder/Non-Responders was observed across the expected eigenvectors (**Fig. S2B**). Furthermore, receiver operating characteristic analysis demonstrated that PC1 scores perfectly discriminated Function-Pain Responder from Non-Responder MSC(M) batches (**Fig. S2C**).

**Fig. 2.**
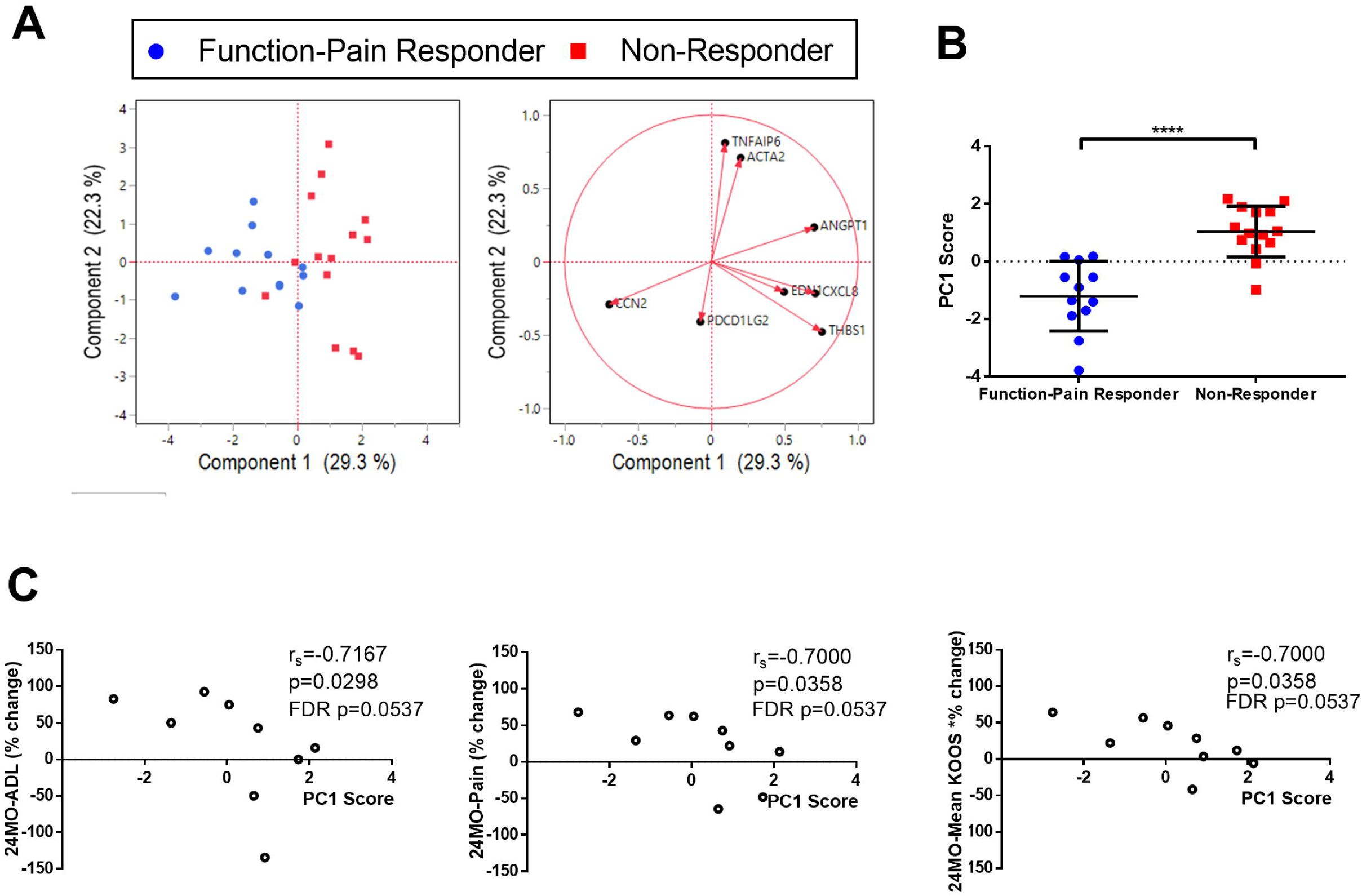
Independent validation that a previously identified gene panel distinguishes clinically effective MSC(M) donors from non-responder MSC(M) donors. A) Principal component (PC) analysis of MSC(M) expression of an immunomodulatory gene panel identified previously for MSC(AT) (*TNFAIP6*, *ACTA2*, *ANGPT1*, *CXCL8*, *EDN1*, *THBS1*, *PDCD1LG2*, *CCN2*; [18]) expressed under licensed conditions. The corresponding loading plot of eigenvectors (right) indicates the relative contribution of each gene to the PC1 and PC2 axes. B) MSC(M) derived from Function-Pain Responders displayed significantly lower PC1 scores relative to non-responders, indicating that the gene panel sensitively distinguishes MSC(M) from donors classified as responders. Student’s t-test, ****p<0.0001. Horizontal lines: group mean; error bars: standard deviation. C) Significant negative correlations between PC1 scores and the percentage change at 24-months relative to baseline for KOOS ADL, pain, symptoms, and mean KOOS. Percent change in KOOS values were made relative to baseline and calculated such that positive values represent improvement. Spearman’s correlation. N=9 MSC(M) donors; n=2-3 replicates/donor; all technical replicates are displayed in A&B; the average of technical replicates for each donor is displayed in C.

PC1 scores, indexed as a composite score for the basal immunomodulatory gene expression profile, additionally showed statistically significant robust negative correlations with improvements in 12-and 24-month mean KOOS and KOOS pain and/or ADL sub-scores at 12- and 24-months (**Fig. 2C**, **Table S1**).

Individual MSC(M) genes, *ANGPT1* and *CCN2* showed significant correlations with 12-and/or 24-month KOOS Pain and ADL subscores and mean KOOS (**Table S2**) that were lost after adjustment for multiple comparisons, suggesting that the multivariate gene expression panel could better distinguish Function-Pain Responder MSC(M) from Non-Responder MSC(M) compared to individual genes.

### *In vitro* immunomodulation by donor MSC(M) batches correlates to patient responder status

We further assessed baseline MSC(M) immunomodulatory potency via functional *in vitro* monocyte polarization using non-donor matched human peripheral blood monocytes (**Fig. 3A**). Discriminant analysis demonstrated distinct clustering and statistically significant separation of monocyte phenotypic profiles between Function-Pain Responder vs. Non-Responder donor MSC(M) batches, with few misclassified data points and a high Entropy R^2^ (**Fig. 3B**, **Table S3**). Co-cultures of monocytes with Function-Pain Responder MSC(M) donor batches showed non-significant upregulation of the inflammation-resolving markers *CD163*, *IL10* and *STAB1,* and reduced expression of *CD86* a*nd CCR7* compared to Non-Responders batches (**Fig. 3B**). The immunosuppressive marker, *CD274*, was significantly upregulated by Non-Responder vs. Function-Pain Responder MSC(M) donor batches. Overall, Function-Pain Responder MSC(M) donor batches modulated *in vitro* monocyte polarization toward a distinct phenotype.

**Fig. 3.**
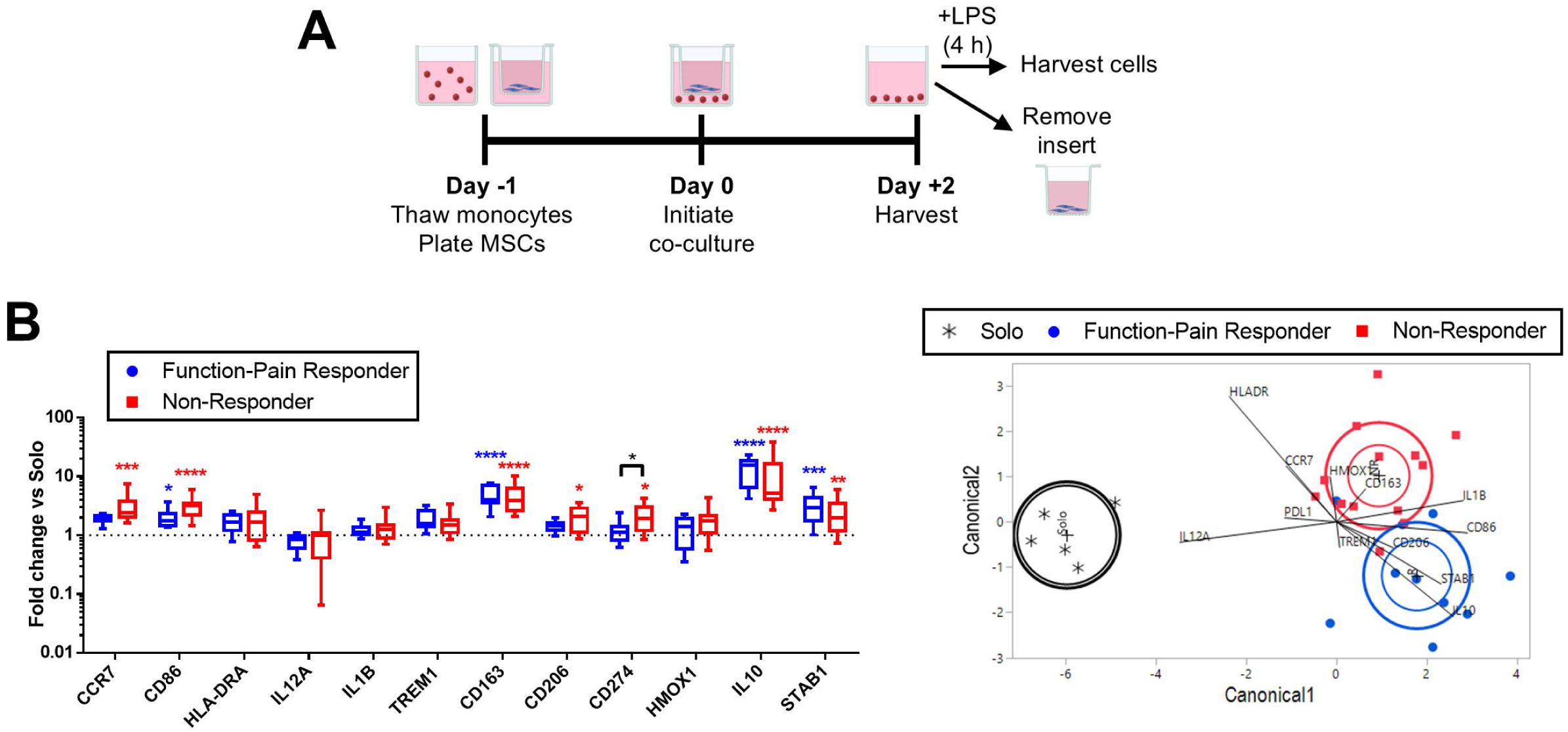
*In vitro* monocyte/macrophage polarization readouts distinguish donor MSC(M) immunomodulatory fitness for Function-Pain Responders versus Non-Responders. A) Schematic of *in vitro* monocyte/macrophage (MΦ) polarization experiment. Indirect co-cultures of MSC(M) with human peripheral blood-derived MΦ was performed for 48 h prior to removal of MSC(M) and addition of lipopolysaccharide (LPS) for 4 h incubation. B) Summary of MΦ gene expression after co-culture with Function-Pain Responder MSC(M). Box-and-whisker plots (left) display changes in gene expression relative to MΦ cultured alone (Solo; dotted line). Horizontal line: median; hinges: first and third quartiles; whiskers: range. Two-way ANOVA, Tukey’s post-hoc test. *p<0.05, **p<0.01, ***p<0.001, ****p<0.0001 relative to Solo condition or to groups indicated by brackets. Discriminant canonical plots (right) of -ddCt values relative to Solo condition. Inner ellipses indicate 95% confidence for the mean of each group; outer ellipses indicate the normal region estimated to contain 50% of the population for each group.

Using discriminant analysis, we noted that re-classification of MSC(M) from patient 10 (classified as a 12-month Non-Responder, but a 24-month Responder) as a Responder improved model fit with fewer misclassified data points and higher Entropy R^2^ relative to the model for 12-month Function-Pain Responder classification (**Table S3**, **Fig. S3**). This classification system also showed a more distinct pro-resolving monocyte phenotype mediated by Responder MSC(M) batches as characterized by significantly higher *IL10* and *STAB1*, and lower *CD86* vs Non-Responders (**Fig. S3**).

### Specific microRNAs distinguish responder donor MSC(M) batches

MicroRNA-sequencing was used to probe differences in MSC(M) donor batches after synovial fluid stimulation. PCA demonstrated clustering by donor batch and responder group rather than by synovial fluid stimulation (**Fig. 4A**). Fourteen significantly differentially expressed microRNAs were identified between Function-Pain Responder vs. Non-Responder MSC(M) donor batches after correction for multiple comparisons; 8 microRNAs were upregulated, and 6 were downregulated (**Fig. 4B**, **Table S4**). Gene targets related to cell cycle (GTPases), phosphoinositide 3-kinase (PI3K), mitogen-activated protein kinase (MAPK), transforming growth factor beta (TGF-β), MyD88, circadian clock genes, SMAD proteins, endocytosis, and other pathways were identified (**Fig. 4C,D**). Gene ontology analysis identified biological processes linked to neuronal processes, TGFβ signaling, cell growth, and Wnt signaling (**Fig. 4E,F**).

**Fig. 4.**
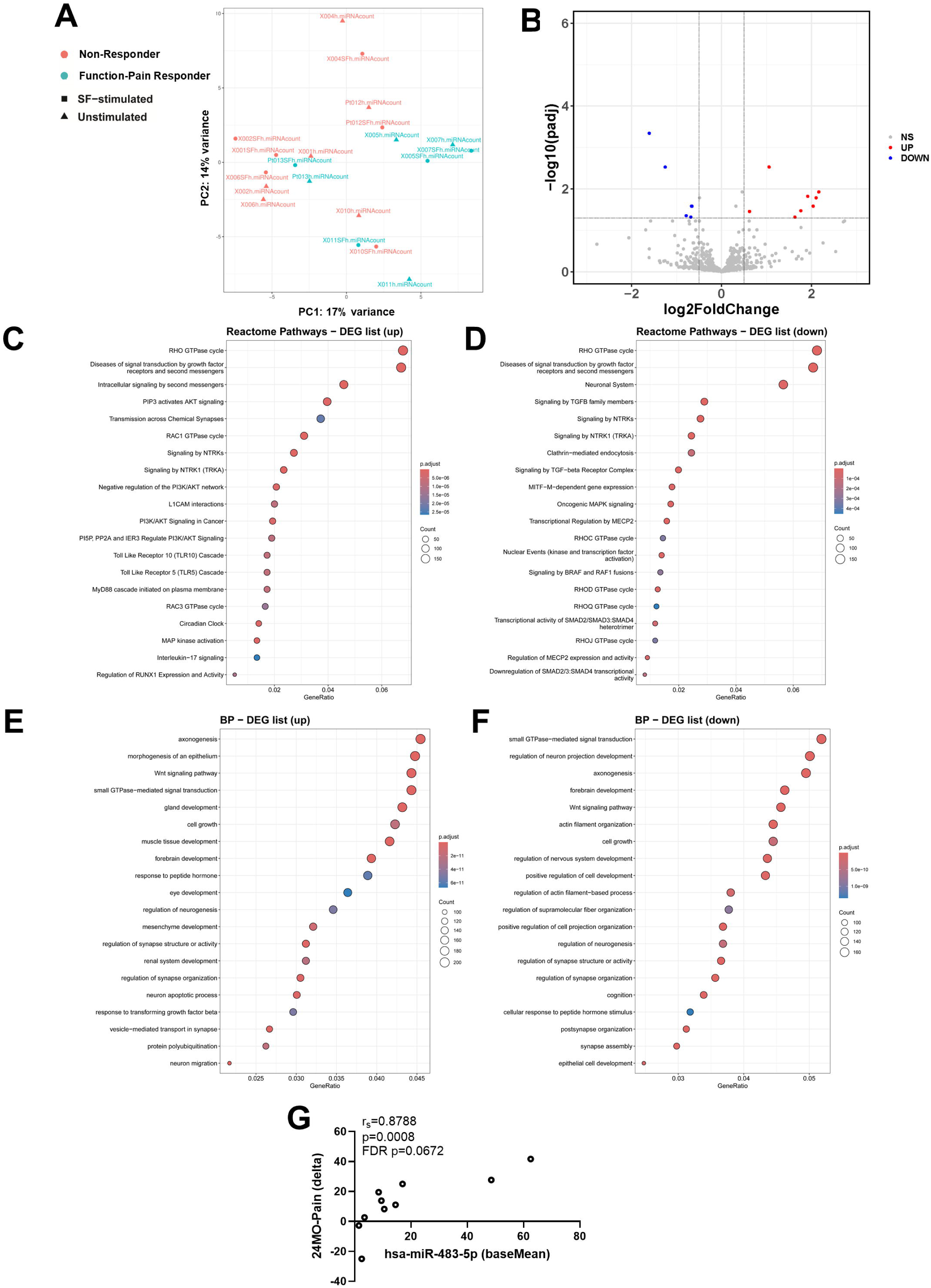
microRNA-sequencing reveals microRNA signatures in Function-Pain Responder donor MSC(M). A) Principal component analysis of microRNA expression profiles demonstrate clustering in Function-Pain Responders versus Non-Responders. MSC(M) samples cluster by donor and responder status rather than by synovial fluid (SF) stimulation. B) Volcano plot displays 14 significantly differentially expressed microRNAs in MSC(M) derived from donors classified as Function-Pain Responders versus Non-Responders. C&D) Top 20 pathways identified through Reactome analysis of predicted gene targets for upregulated (C) and downregulated (D) microRNAs. E&F) Top 20 pathways identified through Gene Ontology (Biological Processes) analysis of predicted gene targets for significantly upregulated (E) and downregulated (F) microRNAs. G) Significant correlation between hsa-miR-483-5p and the delta at 24-months relative to baseline for KOOS pain. Delta change in KOOS values were made relative to baseline and calculated such that positive values represent improvement. Spearman’s correlation. N=10 MSC(M) donors (data points represent each MSC(M) donor treated with or without synovial fluid).

We queried our MSC(M) gene panel against predicted gene targets of the differentially expressed microRNAs and identified *ANGPT1* as a predicted target of hsa-miR-338-5p (a microRNA upregulated in Function-Pain Responders), aligning with the trend toward reduced *ANGPT1* observed in Function-Pain Responders. However, we also identified *ANGPT1* as a predicted target of hsa-miR-210-3p (a microRNA downregulated in Function-Pain Responders), reflective of complex interactions between microRNAs on gene networks, including indirect effects. Limited correlations between the 14 differentially expressed microRNAs with the curated gene panel were identified, but significance was lost after correction for multiple comparisons (**Table S5**), suggesting putative indirect modulation of select MSC(M) genes by differentially expressed microRNAs that needs to be confirmed in larger datasets. Of the significantly differentially expressed microRNAs, hsa-miR-483-5p positively correlated with improvements in 24-month KOOS pain (**Fig 4G**; **Table S6**). Differentially expressed microRNAs provides further characterization and potential mechanistic insights into MSC(M) therapeutic effects in KOA patients.

### Baseline clinical phenotype correlates with 12- and 24-month improvements

We conducted exploratory analysis between baseline inverted KOOS subscales with 12-month responder status (**Fig. 5**). Function-Pain Responders displayed elevated mean KOOS and KOOS subscales at baseline compared to Non-Responders, supportive of more severe baseline clinical presentations in directing MSC responsiveness, albeit without significance (**Fig. 5A**). Baseline MRI WORMS and synovitis scores were not significantly different between Function-Pain Responders and Non-Responders (**Fig. 5B**).

**Fig. 5.**
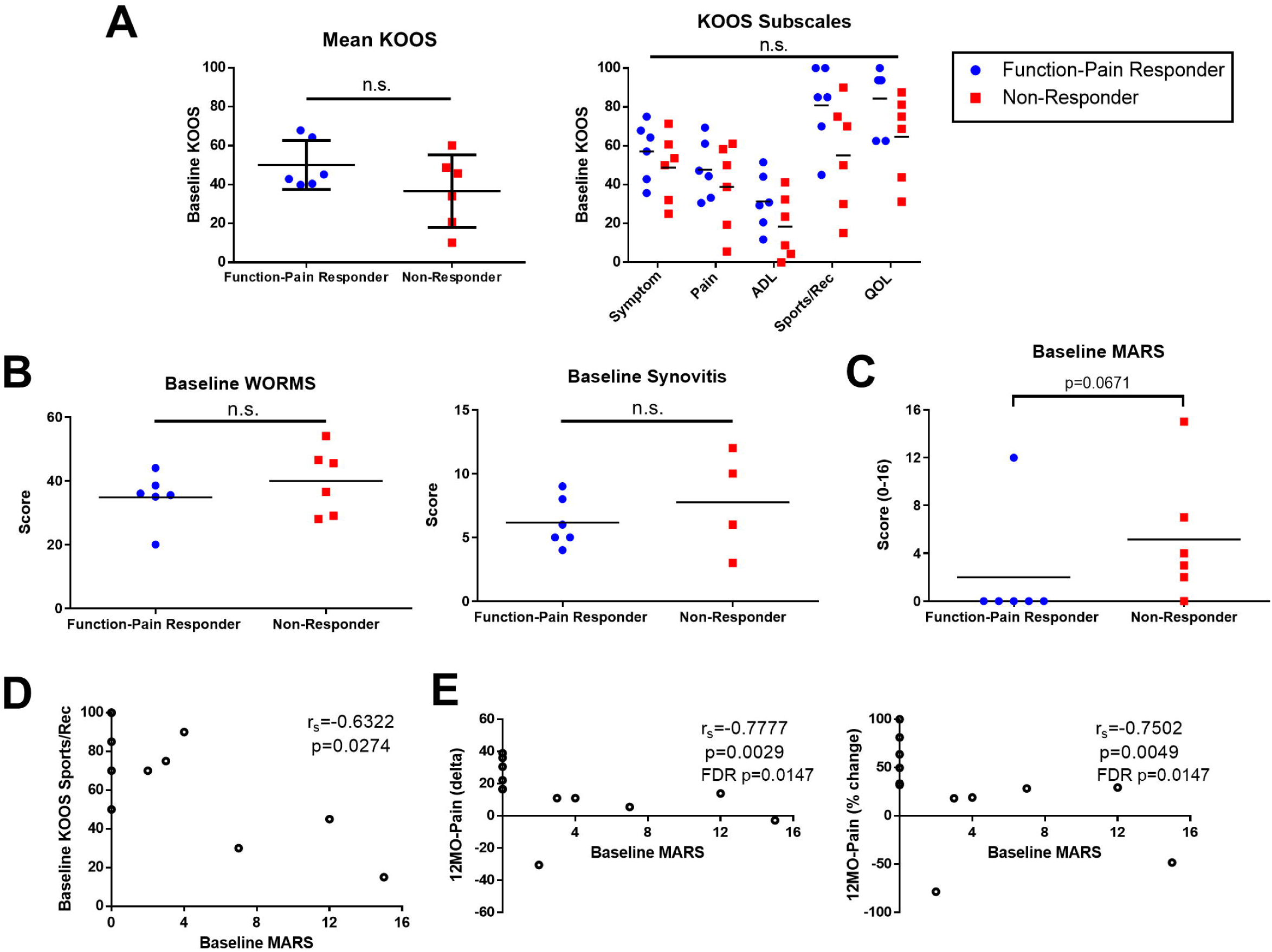
More severe KOOS and lower MARS physical activity scores correlate to improved patient-reported outcomes at 12- and 24-months follow-up. A) Patients categorized according to Function-Pain Responders displayed a trend toward more severe KOOS at baseline relative to Non-Responders (non-significant). B) Baseline MRI WORMS and Synovitis scores were not significantly different in Function-Pain Responders vs Non-Responders. Welch’s t test. C) Baseline MARS scores trended higher in Non-Responders vs Function-Pain Responders. Mann-Whitney test. D&E) Baseline MARS significantly correlated to baseline Sports/Rec, as well as both delta and percent change in KOOS Pain at 12 months relative to baseline. KOOS scales were inverted such that higher scores represent greater knee-related issues; delta and percent change in KOOS values were made relative to baseline and calculated such that positive values represent improvement. N=12 patients. Student’s t-test (for baseline Mean KOOS), Two-way ANOVA, Sidak’s post-hoc test (for baseline KOOS Subscales), Spearman’s correlation. n.s.; non-significant. ADL: function in daily living; Sports/Rec: function in sport and recreation; QOL: knee-related quality of life; Pt.: patient ID; MO: month; KOOS: knee injury and osteoarthritis outcome score; MARS: Marx Activity Rating Scale.

Next, we analyzed correlations between the baseline scores for all KOOS subscales with changes in these scores at 12- and 24-months (**Table S7**). Only baseline Sports/Rec scores positively correlated with improvements in KOOS ADL at 12- and 24-months; however, significance was lost after correction for multiple comparisons.

Given the findings on KOOS Sports/Rec, we probed the relationship with physical activity levels using the MARS scores [26]. A trend (p<0.1) toward higher baseline MARS scores (i.e., more frequent physical activity) was observed in Non-Responders vs Function-Pain Responders (**Fig. 5C**). Higher MARS scores were significantly associated with lower (i.e., less severe) baseline KOOS Sports/Rec scores (**Fig. 5D**). Interestingly, lower baseline MARS scores significantly correlated with improvements in the 12-month KOOS Pain sub-score (**Fig. 5E**, **Table S8**).

### Baseline MRI scores did not correlate with responder status

We had previously reported no significant improvements in MRI WORMS or synovitis scores, calculated relative to baseline based on blinded analysis by two radiologists [8]. Baseline MRI WORMS and synovitis scores did not correlate with responder status (**Table 1**) or to improvements in 12- and 24-month KOOS (**Tables S9, S10**).

### Baseline synovial fluid biomarkers do not correlate with 6, 12- and 24-month patient improvements

The levels of baseline inflammatory, angiogenic, metabolic and cartilage extracellular matrix-associated biomarkers in synovial fluid from a subset of nine patients were previously reported [8]; here we analyzed correlations with 6, 12- and 24-month KOOS improvements (**Tables S11, S12**). There were **<u>negative</u>** correlations for baseline HGF and VEGFA with greater improvements in KOOS Pain subscores at 12-months, but significance was lost after adjustment for multiple comparisons (**Table S11**). No significant correlations were seen between other local biomarkers including TIMP1, resistin, adipsin, leptin, adiponectin, soluble (s)CD14, sCD163, MMP1, MMP9, IL6, IL8, CXCL1, CCL2, CX3CL1 with mean KOOS and KOOS pain or function sub-scores for all timepoints. [8].

### Exploratory analysis of baseline synovial fluid immune cell phenotypes

As additional explorations of KOA patient molecular endotype, we analyzed synovial fluid immune cells using previous data [8]. While this analysis was feasible for a subset of five patients due to limited synovial fluid volumes, our data provides proof-of-concept for probing baseline levels of CD14^+^CD16^+^ and CD14^+^CD163^+^ monocyte/macrophages (MΦs) (**Fig. S4A**) as well as CD3^+^CD4^+^CD69^+^ and CD3^+^CD4^+^CD25^+^ T helper cells (**Fig. S4B**) as putative immunophenotypes of interest. Higher baseline frequencies of Mφs and activated T cells were present in Non-Responders, although without statistical testing given the limited sampling.

### Baseline systemic biomarker levels do not correlate with patient improvements

Systemic baseline inflammatory biomarkers were measured in the present study (sCD163; CRP; S100A8/A9, TNFα; IL-6) alongside previous data on systemic biomarkers of cartilage breakdown [8] (**Table S13, S14**). Higher levels of baseline plasma TNFα significantly positively correlated to percentage changes in KOOS Pain subscores at 6 months (**Fig. S5**, **Table S14**), but this significance was lost with multiple comparison corrections. Thus, the selected panel of local and systemic biomarkers of different categories showed limited capacity to correlate with patient-reported outcomes.

## DISCUSSION

Statistically powered clinical trials using MSCs have reported conflicting results in KOA patients [5,12]. To address underlying factors that drive heterogeneous responsiveness to MSCs, we analyzed our test clinical dataset [8] and identified putative autologous MSC(M) donor batch characteristics and responsive KOA patient features that correlated with improvements in PROMs at 12- and 24-months. We anticipate that this combinatorial analysis will serve as an integrated algorithm that can be tested in prospective KOA clinical trials (and retrospectively on existing datasets) to i) screen potent donor/batches of MSCs; and ii) to identify characteristics of responsive KOA recipients. Our data suggests that autologous MSC(M) donor batches with high basal immunomodulatory fitness as measured by reduced levels of angiogenic genes, distinct monocyte modulatory activity and a distinct signature of microRNAs will be more clinically effective. Our analysis of responsive KOA patients was more exploratory given the limited dataset. We identified trends in KOA patients with reduced knee functionality associated with sports and recreation activities, more severe baseline KOOS and reduced physical activity levels, who were more responsive to autologous MSC(M) injections. Biomarker analysis did not show significant correlations albeit trends pointing towards reduced baseline levels of soluble angiogenic factors in the local joint environment that need to be further investigated in larger clinical datasets with matched placebo controls. We provide a summary of baseline MSC(M) characteristics and baseline patient clinical phenotype/molecular endotype features that correlated to improvements in KOOS in **Table 2**.

**Table 2.** Summary of MSC(M) basal fitness, baseline clinical phenotype and molecular endotype features that correlate to patient responder status in KOA.

|  |  | **Function-Pain Responder Characteristics** |
| --- | --- | --- |
| **MSC(M)**<br>**Basal**<br>**Fitness** | **Gene**<br>**expression** | Distinct gene expression profile (Lower PC1 score driven by higher *CCN2* and lower *THBS1*, *CXCL8*, *ANGPT1*, etc.) |
|  | **Monocyte/**<br>**macrophage**<br>**polarization** | Responder MSC(M) batches induce a distinct monocyte/macrophage gene expression profile |
|  | **microRNA**<br>**expression** | 14 significantly differentially expressed microRNAs vs Non-Responders |
| **Baseline**<br>**clinical**<br>**phenotype** | **KOOS** | Trend toward more severe KOOS at baseline (non-significant) |
|  | **MARS** | Lower physical activity level at baseline |
| **Baseline**<br>**molecular**<br>**endotype** | **Local**<br>**biomarkers** | Lower baseline levels of HGF and VEGFA (non-significant) |
| | **Systemic**<br>**biomarkers** | Higher baseline levels of TNFα in plasma (non-significant) |

We used a previous clinical trial [8] with twelve manufactured, patient-specific lots of MSC(M), which was data-rich with multiple baseline biomarkers, and clinical readouts and offered an excellent test dataset to study autologous MSC(M) batch variations. We provided longitudinal 12- and 24-month data and are the first to report long-term 8–10-year clinical follow-up. Function-Pain Responders showed significant improvements in PROMs at 12- and 24- months; 50% of patients were classified as having clinically meaningful improvements in both pain and function at 12-months, and 42% of patients sustained these clinically meaningful improvements out to 24-months. We note that 3/6 Function-Pain Responders and 1/5 Non-Responders (one lost to follow-up) received no additional interventions over 8-10 years. In terms of injections over the intervening years; 1/6 Function-Pain Responders received injections vs. 3/5 Non-Responders. Lastly, 2/6 Function-Pain Responders vs. 1/5 Non-Responders received or were awaiting a total knee replacement (TKR). Given the extended lag in the follow-up, we cannot conclude that a single autologous MSC(M) injection delayed knee replacement or reduced additional injections.

Autologous MSC(M) induced both function and pain improvements within 24 months. Although immunomodulatory therapies may be expected to induce greater analgesic effects [27], MSCs act through multimodal mechanisms[28]. Notably, our analysis suggesting that patients with more severe KOA were more responsive to MSC(M) treatment aligns with findings in acute graft-versus-host disease [29], myocardial infarction [30] and ulcerative colitis [31], where more severe disease patients responded better to MSC treatments. Function-Pain Responders showed trending correlations to higher overall disease severity and significantly lower baseline MARS physical activity scores. Patients with higher baseline MARS scores had lesser improvements, suggesting that patients who persisted through pain in physical activity may have exacerbated disease progression.

We focused on basal immunomodulatory properties of MSC(M) and their ability to modulate monocytes, given their importance as a clinically relevant mechanistic target in KOA [4,7,8,32]. Monocyte/macrophage roles in driving synovitis and OA progression has previously been shown [8,33–35]. We measured expression of a previously reported gene panel that correlated to *in vitro* functional activity [18], and determined that the gene panel significantly discriminated between Function-Pain Responder vs. Non-Responders autologous MSC(M) donor batches. Dimension-reduced PC1 scores capturing multivariable genes were strongly and significantly correlated inversely to patient improvements at 12- and 24-months and were thus more reliable than individual genes. Notably, this panel included predominantly angiogenic markers that were previously inversely correlated to monocyte polarization toward inflammation-resolving phenotypes [18], supportive of inverse relationships between angiogenic and immunomodulatory fitness of MSC(M) [36,37]. The genes identified for MSC(AT) were valid for MSC(M) as well, speaking to their broad utility to serve as MSC CQAs [7]. These data support our hypothesis that MSCs with high basal immunomodulatory fitness produce greater improvements in knee function. Nonetheless, the use of autologous MSCs [7,8] adds a layer of complexity, given that systemic factors such as age, BMI, and other health conditions were dually accounted for in both basal fitness of the cells [38,39] and baseline patient disease status [40].

Function-Pain Responder autologous MSC(M) donor batches modulated CD14^+^ monocytes to more pro-resolving subtypes *in vitro*, although there was also upregulation of pro-inflammatory markers. There is growing recognition that monocytes are highly plastic, and multivariable characterization provides a more nuanced recognition of their mixed and dynamic transcriptomic and phenotypic profiles [41–43]. Reliance on dichotomized gene expression markers is not indicative of the plurality of monocyte functionality [44]; rather induction of mixed monocyte phenotypes by MSCs is important for achieving therapeutic effects [45]. Importantly, we have shown that CD14^+^ monocytes in OA synovial fluid, despite exhibiting a mixed phenotype when co-cultured with MSC(M), are functionally reparative [46].

Function- Pain Responder MSC(M) donor batches had a significantly different microRNA signature compared to Non-Responders. Pathways related to inflammation, fibrosis, angiogenesis and cell cycle, relevant to KOA pathophysiology were identified. We also identified enrichment of pathways involved in cell cycle. Our analysis identified enrichment of several neuronal processes, which given the role of innervation and neuronal signaling in joint pain in KOA [47] implicates these microRNAs in mediating analgesic effects of MSC(M). The functionality of differentially expressed microRNAs requires further study but serves as additional CQAs for identifying basally fit MSC(M).

Using a curated panel of inflammatory, angiogenic, and metabolic proteins and proteases, we could not discern a signature of local biomarkers that definitively identified responsive KOA patients to MSC(M). Responders displayed putatively lower local baseline levels of angiogenic biomarkers, HGF and VEGFA, but this lost significance after multiple comparisons correction. Systemically, there was limited short-term positive correlations between TNFα (plasma) levels and PROM improvements, albeit significance was again lost after multiple comparisons correction. Baseline serum CRP levels, an inflammatory biomarker in multiple inflammatory conditions [48] and cartilage breakdown markers (shown to correlate with worsening KOA pain [49]) failed to correlate with improvements in KOOS at all time points. Overall, our analyses failed to show significant correlations to MSC(M) treatment responsiveness with any of the biomarkers tested. However, a trend towards a mixed profile with increased pro-inflammatory systemic TNFα but reduced local angiogenic proteins (HGF and VEGF) emerged, reminiscent of mixed biomarker profile in rheumatoid arthritis patients, responsive to allogenic MSCs [50]. This mixed local vs. systemic molecular endotype profile needs to be further tested in larger datasets.

Identifying molecular endotypes of OA has proven to be extremely challenging, and multi-consortium efforts have not been able to definitively identify molecular endotypes when profiling local synovial fluid, even with large, covariate-adjusted datasets [51]. This heterogeneity in OA patients is a key reason for failure of multiple drugs in late-phase, powered clinical trials [52,53]. The absence of stratification biomarkers to identify OA theratypes for which specific treatments can be tailored to specific clinical phenotypes and endotypes has been identified as a critical gap impeding the development of disease modifying drugs in the field [54]. Although systemic biomarkers of inflammation and/or cartilage degradation are beginning to emerge to cluster patients [55], their utility in defining patient responsiveness to different therapies remains untested. In our hands, many of the previously reported inflammatory and cartilage degradation biomarkers [8,55] did not correlate with improvements in patient-reported outcomes to MSC(M) treatments.

Limitations of this study include small sample size, which allowed us to perform an exploratory but not definitive analysis. Indeed, baseline clinical phenotype and local biomarkers indicative of reduced angiogenic and inflammatory molecular endotype lost significance when corrected for multiple comparisons, likely due to overcorrection arising from the small sample size.

We also did not adjust for sex, age, BMI, MSC dose or OA stage given small sub-group size within our limited dataset; we justified this based on absence of correlation of these factors with Function-Pain Responder/Non-Responder status. Nonetheless, these adjustments are important and should be considered in future analysis and application of our proposed algorithms in larger clinical trial datasets. Even with this, we noted trends in dose response; three of four patients in the higher MSC(M) autologous dose group (50×10^6^ MSCs) were classified as Function-Pain Responders, aligning with our original analysis [8] and review [4]. Dose dependencies are not always clearly delineated for MSC therapies that work by interacting with variable host immune effectors including monocytes/macrophages [4,8,33]. Thus, MSC effect depends equally on baseline patient immune and inflammatory status, which has been difficult to untangle [56]. Indeed, in dose-response studies of allogenic MSC in KOA, lower doses have been favoured [57,58], further exemplifying the complex relationship between MSC dose and patient baseline status.

Another key limitation was the absence of a control group in the original clinical trial [8], although effects were measured at multiple timepoints including 3, 6, 12- and 24-months. KOA is a progressive condition [59], with approximately 30% of patients displaying worsening outcomes over two years [60]. While placebo effects cannot be ruled out, less than 30% of patients reported worsening outcomes, potentially speaking to a treatment effect. Indeed, analysis of non-responders showed that even patients within this category did not show any significant worsening in PROMs over 24-months.

Despite these limitations, the multivariate framework we proposed of investigating potency features of MSC investigational products combined with clinical and biomarker data to analyze patient phenotypes and endotypes is a useful method to test and stratify OA patients into theratypes, suited for MSC therapies [54].

Donor batches of MSC(M) that resulted in positive clinical outcomes differed significantly in specific measurable attributes. These quantitative, multivariate, and functionally relevant attributes can define future CQAs for MSC(M) that can be evaluated to strategically design KOA trials to ensure higher success rates of MSC injection. Additionally, exploratory baseline clinical features and biomarkers identified in responsive KOA patients to MSC(M) injections form the basis for further exploration in larger clinical datasets to verify their utility.

## METHODS

### Clinical trial and patient data

KOA patients (Kellgren Lawrence (KL) grade 3-4) were recruited to a nonrandomized, open=:Jlabel, dose=:Jescalation, Health Canada-authorized phase I/IIa clinical trial as reported [8] (NCT02351011). Ethics approvals including for long-term follow up were granted (REB 14–7909;20-5066;24-5463) [8]. Complete clinical follow-up was performed 24-months after MSC(M) injection for all patients. Additional follow-up at 8-10 years was obtained for 10/12 patients; 2 patients were lost to follow-up. Calculation of patient-reported outcome measures (PROMs), including the inverted Knee Injury and Osteoarthritis Outcome Score (KOOS) and the Marx Activity Rating Scale (MARS), is described in **Suppl. File 1**. Contrast-enhanced MRI analysis at baseline, 6 and 12-months were conducted [8] along with whole organ MRI scores (WORMS) and MRI synovitis scores calculated as described [61].

### Responder vs. Non-Responder Criteria

Patient clinical responder status was based on the Outcome Measures in Rheumatology (OMERACT) and Osteoarthritis Research Society International (OARSI) Standing Committee for Clinical Trials Response Criteria Initiative (OMERACT-OARSI criteria), based on data from large, powered placebo-controlled drug trials for hip and KOA [23]. These criteria categorize patients as responders according to the following: i) substantial improvement in pain or in function ≥50% and absolute change (delta) ≥ 20; OR, ii) significant improvement (≥20% and absolute change (delta) ≥ 10) in at least **two** of the following: a) pain, b) function, c) patient global assessment; the latter was not measured in our trial. We categorized patient responder status using the twelve-month follow-up data based on changes in ***both*** KOOS function (activities of daily living (ADL) subscale) and KOOS Pain, directly applying OMERACT-OARSI criteria thresholds for function and pain scores of delta value ≥10 AND percentage change ≥20% [23]. At the 24-month time point, one patient (#3) exceeded the substantial threshold of ≥50% and absolute change (delta) ≥ 20 in KOOS ADL without ≥20% and absolute change (delta) ≥ 10 changes in KOOS Pain; the patient is therefore still considered a Responder according to OMERACT-OARSI criteria. Thus, our classification criteria follows the stringent requirement for significant, clinically-meaningful improvements of at least 10 points [62,63] based on pain and function, two of three recommended metrics. The validated KOOS instrument measures responses to interventions in KOA with high test-retest reliability and improved sensitivity in pain and function scoring relative to the Western Ontario and McMaster Universities Osteoarthritis Index (WOMAC) scoring system [23,64]. Baseline demographics, radiographic KL-grade, magnetic resonance imaging (MRI) total cartilage volume (WORMS) [61] and synovitis [65] scores are provided for Function-Pain Responder vs. Non-Responders (**Table 1**). MSC(M) culture and characterization, including gene expression (**Table S15**), microRNA-sequencing, and *in vitro* monocyte polarization are described in **Suppl. File 1**.

### Biomarker measurements

Levels of curated biomarkers in baseline serum (**Table S16**) were measured previously [8]; updated analyses and experimental questions were applied to the dataset in the present work. Additional inflammatory biomarkers were measured in baseline biobanked patient serum and plasma (analytes specified in **Table S16**). Synovial fluid collection was feasible for only nine patients. Exploratory immune cell profiling was done only on available (five) synovial fluid samples [8].

### Statistical analyses

GraphPad Prism 6.0 and JMP Pro 17 software were used for statistical analyses and to create plots. Statistical tests are specified in figure and table captions. Spearman’s correlation was selected as a non-parametric statistical test to investigate associations between inverted KOOS changes and other variables, given that the data violated assumptions of normality and homoscedasticity. P values derived from Spearman’s correlation were corrected for multiple comparisons using the Benjamini-Hochberg False Discovery Rate (FDR) method. Given the small sample size, covariate adjustment for sex, age, body mass index (BMI), KL-grade, MSC dose and baseline MRI WORMs and synovitis scores were not performed, given lack of significant correlation between these factors with Function-Pain Responder status (**Table 1**). Given the limited sample size and exploratory nature of analysis, Spearman’s correlations were considered statistically significant when p<0.05 (uncorrected) and FDR<0.1.

## Supporting information

Graphical Abstract

## Acknowledgements

We thank Kim Perry, Mary Nasim, Tamara Wagner, Amanda Weston and the Orthopedic Surgery team for assistance in acquiring donor samples. Thanks to Dr. Mozhgan Rasti who performed qPCR for a subset of experiments, as well as Dr. Mohit Kapoor, Dr. Starlee Lively, Dr. Mehdi Layeghifard, and Pratibha Potla for assistance with microRNA-sequencing and analysis. Schematics for figures were created using BioRender.com.

This research was funded by Canadian Institutes of Health Research (CIHR) (PJT-166089) and Natural Sciences and Engineering Research Council of Canada (NSERC) (RGPIN-2018-05737) grants awarded to SV. The work is in part supported by the Schroeder Arthritis Institute via the Toronto General and Western Hospital Foundation (University Health Network). Salary support for KR was provided by a Natural Sciences and Engineering Research Council (NSERC) Canada Graduate Scholarship. Salary support for RR was provided by the Arthritis Society (TPF-19-0537). AP is supported by an award from Arthritis Society Canada (STAR-20-0000000012).

## Author contributions

Kevin P. Robb: Conception & design, collection and/or assembly of data, data analysis and interpretation, manuscript writing, final approval of manuscript

Razieh Rabani: Conception & design, collection and/or assembly of data, data analysis and interpretation

Shabana Vohra: data analysis and interpretation

Shoba Singh: administrative support, collection and/or assembly of data

Oreoluwa Kolade: data analysis and interpretation

Jaskarndip Chahal: collection and/or assembly of data, provision of study materials or patients

Julie Audet: data analysis and interpretation

Anthony V. Perruccio: data analysis and interpretation Ali Naragahi: data analysis

Rajiv Gandhi: financial support, provision of study materials or patients Osvaldo Espin-Garia: data analysis

Sowmya Viswanathan: Conception & design, manuscript writing, final approval of manuscript

