## Supplementary File for "Responders vs. non-responders to mesenchymal stromal cells in knee osteoarthritis patients: mechanistic correlates of donor cell attributes and putative patient features"

Supplementary File 1

SUPPLEMENTARY MATERIALS & METHODS

**MSC(M) isolation and culture**

Biobanked autologous MSC(M) were retained in liquid nitrogen as individual batches by consent of KOA patients. MSC(M) were thawed in three batches at P3 or P4 and plated directly on 24-well plates for microRNA-seq and gene expression analyses. For monocyte polarization experiments, MSC(M) were thawed in three batches at P4 or P5, expanded for one passage, and then plated for the assay. MSC(M) culture was performed using proliferation medium consisting of low-glucose DMEM (Life Technologies) supplemented with 1% Glutamax (Life Technologies) and 10% fetal bovine serum (Hyclone, GE Healthcare, Chicago, USA) on T75 flasks with media changes performed every 3-4 days, as previously described [1]. MSC(M) were harvested at approximately 80% confluency and plated for subsequent experiments as described below.

**Calculation of Knee Injury and Osteoarthritis Outcome Score**

The Knee Injury and Osteoarthritis Outcome Score (KOOS) was applied as a patient-reported outcome measurement tool [2] and the scale was inverted such that higher scores indicate worse outcomes. Mean KOOS was calculated as an average of all responses to the KOOS questionnaire. Both delta and percentage change KOOS values were calculated using scores collected at follow-up time points relative to baseline to capture absolute changes, as well as changes that account for baseline values, respectively. While delta values correspond to the magnitude of difference relative to baseline, they do not account for differences at baseline; percentage change values do control for baseline scores but can be inflated when baseline scores are low. Both are considered relevant measures of change according to international criteria for evaluating osteoarthritis therapeutics [3] and are therefore included throughout this study. Delta and percentage change values were calculated such that positive values indicated improvement. The Marx Activity Rating Scale (MARS) was also used as a tool measuring levels of physical activity for individuals with knee disorders [4].

**MSC(M) gene expression analysis**

MSC(M) from nine donors were seeded on 24-well plates at 30,000 cells/well and licensed with a cocktail of pro-inflammatory cytokines (IFNγ: 30 ng/mL; TNFα: 10 ng/mL; and IL-1β: 5 ng/mL), as previously described [5]. After 24 hours of culture, Trizol-chloroform extraction was performed to isolate RNA, and cDNA was generated using SuperScript™ IV VILO™ Master Mix (Invitrogen, Waltham, USA). FastStart Universal SYBR Green Master Mix (Roche, Basel, Switzerland) and custom primers (**Table S15**) were used for qPCR on a QuantStudio^TM^ 5 system (ThermoFisher). Results were normalized (delta Ct; ∆Ct) using reference genes (*GAPDH*, *GUSB*) and analyzed as negative ∆Ct such that higher (more positive values) indicate a higher level of expression.

***In vitro* monocyte/macrophage polarization**

Indirect co-culture of unmatched human CD14^+^ peripheral blood-derived monocytes (STEMCELL Technologies, Vancouver, Canada) with biobanked MSC(M) was performed as previously described [5]. Co-culture was performed for 48 h and lipopolysaccharide (LPS) was spiked at a concentration of 2.5 ng/mL. LPS was added to elicit a stronger gene expression readout; given the presence of LPS in synovial fluid of KOA patients [6], there is justification for its inclusion in the assay. After 4 h of LPS incubation, monocytes were collected and RNA extraction, cDNA synthesis, and qPCR was performed as described above. Results were normalized (∆∆Ct) using reference genes (*GAPDH*, *B2M*, *TBP*) and presented as fold-change relative to monocytes cultured without MSC(M) (solo condition).

**microRNA-sequencing on MSC(M) samples**

Biobanked MSC(M) from ten donors were thawed and plated on 24-well plates at 50,000 cells/well and cultured for 72 h in proliferation medium, supplemented with or without synovial fluid. Synovial fluid was collected from unrelated late-stage KOA patients (KL grade 3-4, REB #14–7483), pooled from eight patients, and stored at ‑80°C until use, as previously described [7]. For treated wells, medium was supplemented with 30% (v/v) synovial fluid. After 24 h, RNA extraction was performed using a miRNeasy Mini Kit (Qiagen, Hilden, Germany) and stored at -30°C until analysis.

Libraries of cDNA were prepared using the QIAseq miRNA Library Kit (Qiagen) according to the manufacturer's instructions. The libraries were quantified using a DS-11 Spectrophotometer (DeNovix, Wilmington, USA) and assessed for quality using a high-sensitivity DNA chip on the Bioanalyzer (Agilent, Santa Clara, USA). Two samples (one with synovial fluid stimulation and one unstimulated) per biological donor were sequenced using the NextSeq 550 system (Illumina, San Diego, USA) at the Centre for Arthritis Diagnostic and Therapeutic Innovation (Krembil Research Institute, Toronto, Canada) according to previously published methods [8]. Sequencing data alignment and microRNA counts generation was performed as described previously [8]. Single-end 76-base sequencing was performed to an average depth of 2.03 million (± 0.67 million standard deviation) reads per sample. The percentage of sequence alignment was >40% for all samples. All microRNA-seq data has been deposited in NCBI’s Gene Expression Omnibus accessible through GEO Series accession number GSE249452.

Differential expression analysis of microRNA-Seq data was performed using the DESeq2 algorithm developed by Love *et al*. [9]. MicroRNAs were considered significantly differentially expressed based on adjusted p values <0.05 and absolute fold-change >1.5. Given that there were no significantly different microRNAs between synovial fluid-stimulated and unstimulated MSC(M), the samples were pooled for differential expression analysis of responders/non-responder donors. Pathway Analysis was performed on microRNAs that were significantly differentially expressed in Function-Pain Responder vs Non-Responder MSC(M). We performed gene target prediction of the microRNAs as previously described. Gene Ontology Biological Processes enrichment analysis was performed using enrichGO function in clusterProfiler package (version 4.4.4) and retaining terms with q-value lower than 0.05. Reactome pathway enrichment analysis was performed using the ReactomePA package (v1.40.0 in R) based on the REACTOME pathway database [10]. MicroRNA-sequencing data has been deposited in the Gene Expression Omnibus database (Accession number: GSE249452).

SUPPLEMENTARY FIGURES

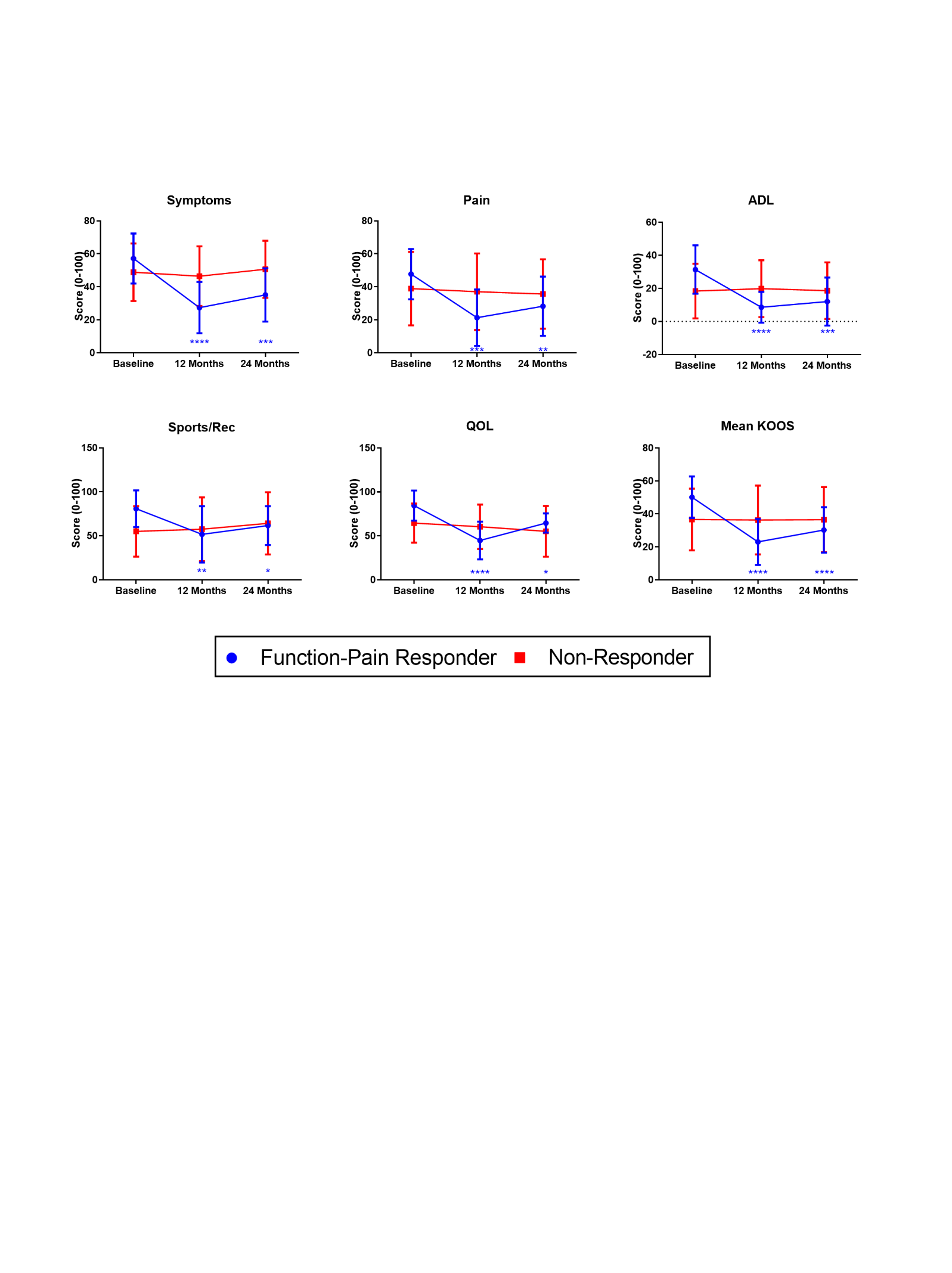

Fig. S1. Function-Pain Responders demonstrate improvements in KOOS outcomes that are maintained out to 24 months follow-up post-injection. Function-Pain Responder and Non-Responder group averages corresponding to the individual patient data plotted in Figure 1A. Two-way repeated measures ANOVA, Dunnett’s multiple comparisons test. *p<0.05, **p<0.01, ***p<0.001, ****p<0.0001; Asterisks indicate significant differences for responder group relative to baseline at the time points indicated. N=12 patients. KOOS scales were inverted such that higher scores represent greater knee-related issues. ADL: function in daily living; Sports/Rec: function in sport and recreation; QOL: knee-related quality of life; KOOS: knee injury and osteoarthritis outcome score.

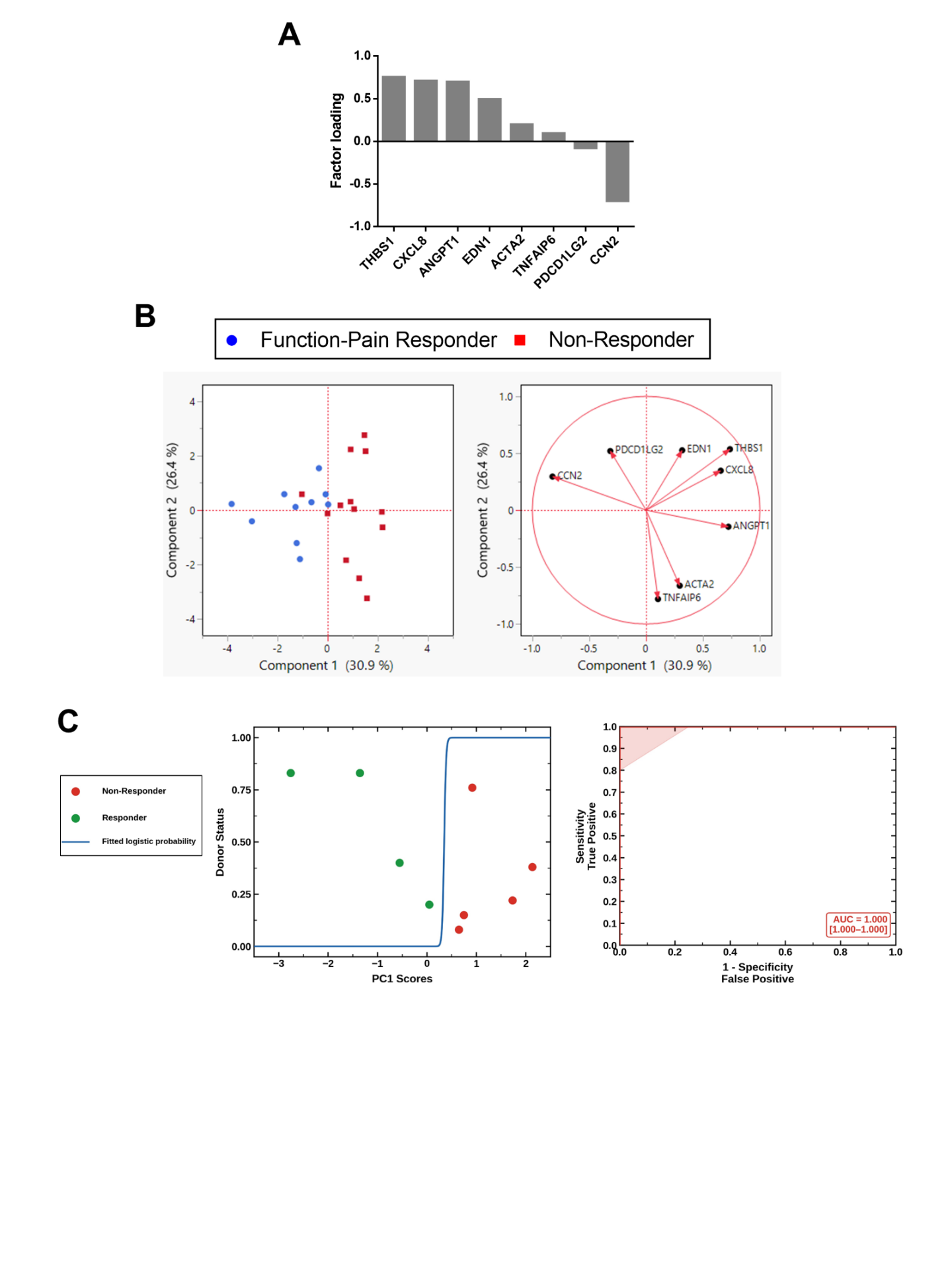

Fig. S2. Putative immunomodulatory CQA genes for MSC(M) derived from Function-Pain Responders. A) Factor loading plot shows the relative contribution of each gene to principal component 1 (PC1) corresponding to the PC analysis in Figure 2A. Loadings close to ±1 indicate a greater relative contribution to PC1. B) Principal component (PC) analysis of MSC(M) expression of an immunomodulatory gene panel expressed under licensed conditions, with outliers removed. The corresponding loading plot of eigenvectors (right) indicates the relative contribution of each gene to the PC1 and PC2 axes. C) Left: Nominal logistic regression of donor response status (Function-Pain Responder vs. Non-Responder) as a function of PC1 scores (n = 9 donors; 7 training, 2 validation). Blue sigmoid curve represents the fitted probability of Non-Responder classification as a continuous function of PC1 score; the decision boundary (p = 0.5) occurs at PC1 ≈ 0.35, above which donors are classified as Non-Responders. The model converged in 18 iterations (Whole Model Test: χ² = 12.37, df = 1, p = 0.0004; McFadden R² = 1.00). Right: Receiver Operating Characteristic (ROC) curve for the logistic model evaluated on the training set (n = 7). The model achieved perfect discrimination (AUC = 1.000 [95% CI: 1.000–1.000]; 20/20 concordant pairs). N=9 MSC(M) donors, n=2-3 replicates/donor.

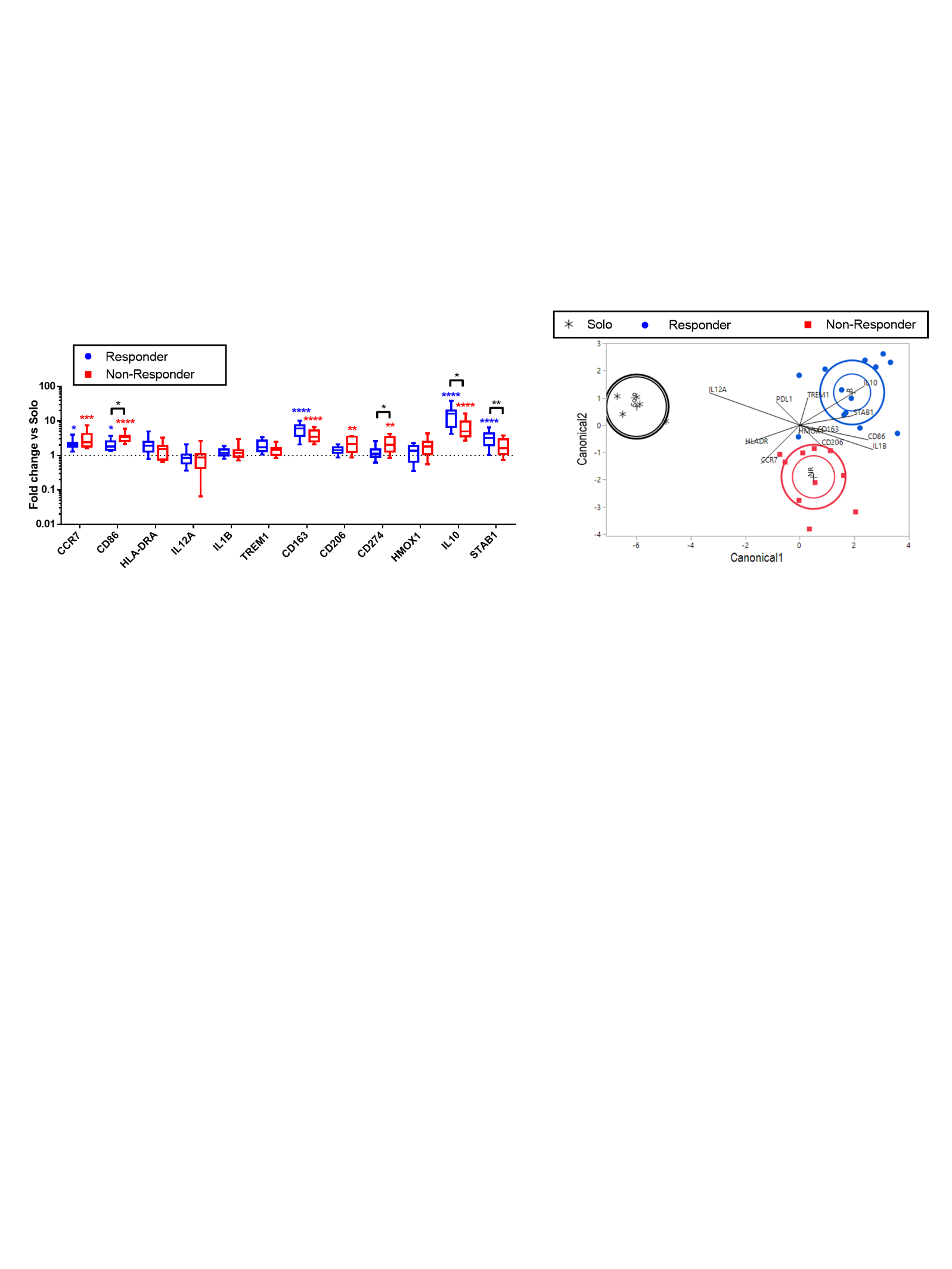

**Fig. S3. *In vitro* monocyte/macrophage polarization readouts distinguish donor MSC(M) immunomodulatory fitness using modified responder classification system.** Summary of MΦ gene expression after co-culture with Responders MSC(M), using patient 10 MSC(M) reclassified as a Responder. Box-and-whisker plots (left) display changes in gene expression relative to MΦ cultured alone (Solo; dotted line). Horizontal line: median; hinges: first and third quartiles; whiskers: range. Two-way ANOVA, Tukey’s post-hoc test. *p<0.05, **p<0.01, ***p<0.001, ****p<0.0001 relative to Solo condition or to groups indicated by brackets. Discriminant canonical plots (right) of -ddCt values relative to Solo condition. Inner ellipses indicate 95% confidence for the mean of each group; outer ellipses indicate the normal region estimated to contain 50% of the population for each group.

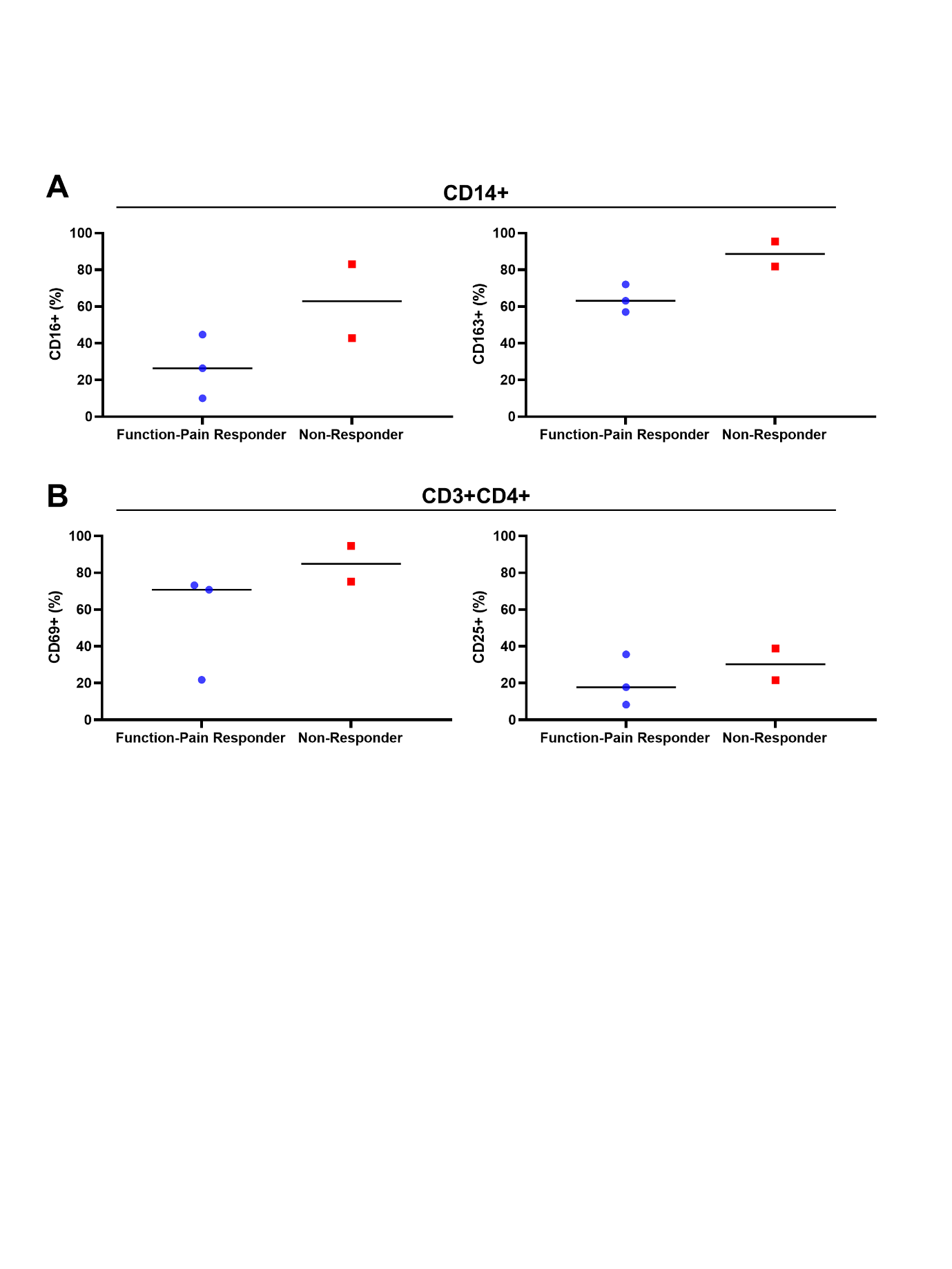

**Fig. S3. Baseline synovial fluid immune cell subpopulations.** A) Levels of CD16+ (left) and CD163+ (right) are expressed as a percentage of total CD14+ MΦ. B) Levels of CD69+ (left) and CD25+ (right) are expressed as a percentage of total CD3+CD4+ T helper cells.

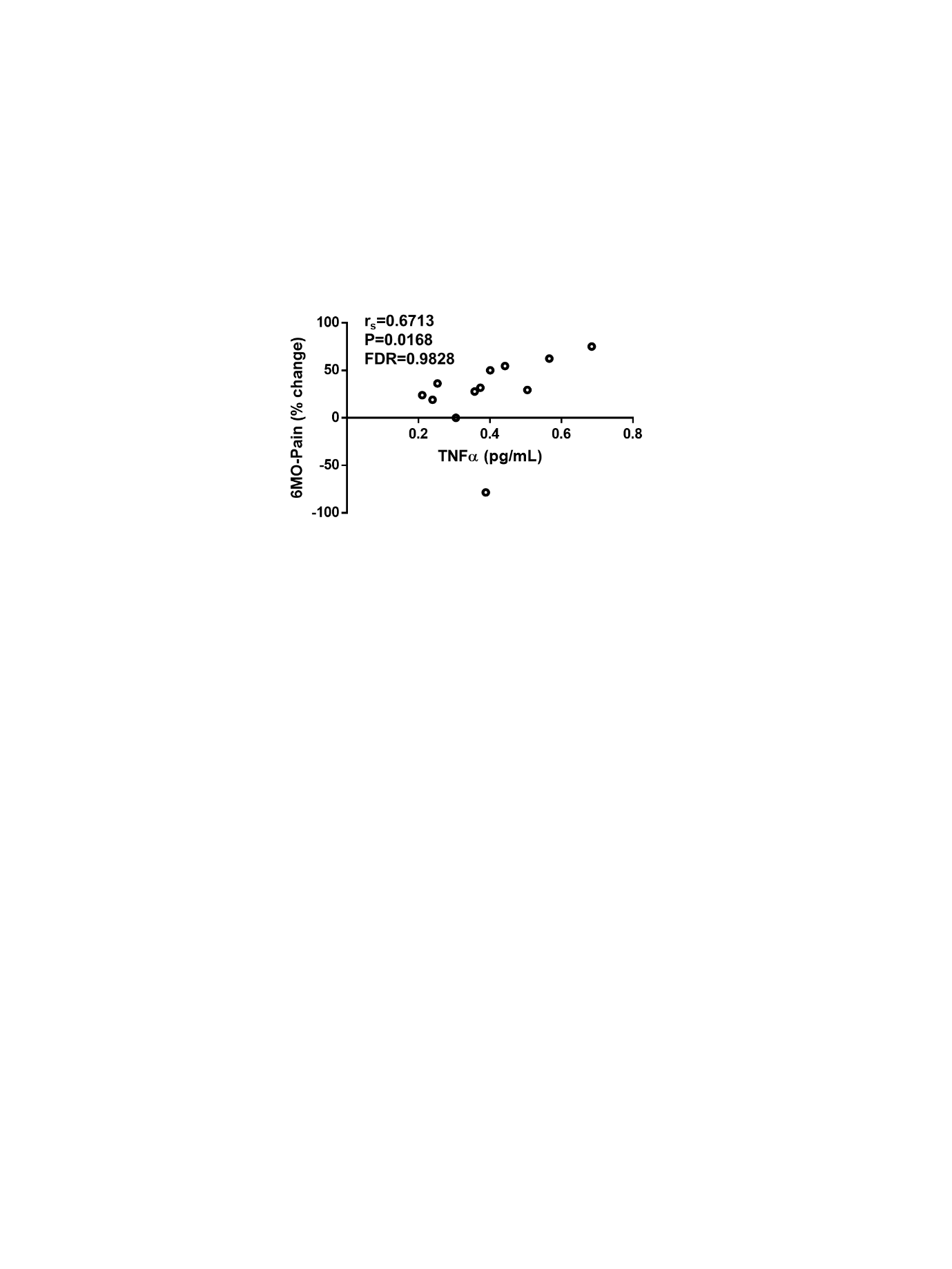

**Fig. S4. Baseline systemic biomarkers correlate with short-term (6 months) but not longer-term patient reported outcome measures.** Higher levels of TNFα measured in plasma at baseline significantly positively correlated to percentage change in KOOS Pain at 6 months relative to baseline. Delta and percent change in KOOS values were made relative to baseline and calculated such that positive values represent improvement. Spearman’s correlation. N=12 patients.

**Table S1: Correlations between PC1 composite score of putative immunomodulatory CQA genes and changes in KOOS (percent change or delta values).** N=9 patients. Significant Spearman’s correlations (*p<0.05, FDR<0.1) are bolded. ADL: function in daily living; KOOS: knee injury and osteoarthritis outcome score; FDR: false discovery rate; PC1: principal component 1.

|  | | | **12 Month** | | | **24 Month** | | |
| --- | --- | --- | --- | --- | --- | --- | --- | --- |
| **PC Score** | **Follow-up Score** | **Difference calculation** | **Spearman's rho** | **p value** | **FDR p value** | **Spearman's rho** | **p value** | **FDR p value** |
| PC1 | Pain | Percent change | -0.5333 | 0.1392 | 0.1392 | **-0.7** | ***0.0358** | **0.0537** |
|  |  | Delta | -0.65 | 0.0581 | 0.0805 | -0.6333 | 0.0671 | 0.0671 |
|  | ADL | Percent change | **-0.7448** | ***0.0213** | **0.0639** | **-0.7167** | ***0.0298** | **0.0537** |
|  |  | Delta | -0.6444 | 0.061 | 0.0805 | -0.6333 | 0.0671 | 0.0671 |
|  | Mean KOOS | Percent change | -0.6333 | 0.0671 | 0.0805 | **-0.7** | ***0.0358** | **0.0537** |
|  |  | Delta | **-0.7667** | ***0.0159** | **0.0639** | **-0.7113** | ***0.0317** | **0.0537** |

**Table S2. Correlations between individual putative immunomodulatory CQA genes and changes in KOOS (percent change or delta values).** N=9 patients. Correlations were corrected for multiple comparisons; given exploratory nature of this analysis, correlations that were statistically significant without adjustment for multiple comparisons correction (*p<0.05, FDR>0.1) are marked with an asterisk. Spearman’s correlation. ADL: function in daily living; KOOS: knee injury and osteoarthritis outcome score; FDR: false discovery rate.

|  | | | **12 Month** | | | **24 Month** | | |
| --- | --- | --- | --- | --- | --- | --- | --- | --- |
| **Gene/** | **Follow-up Score** | **Difference calculation** | **Spearman's rho** | **p value** | **FDR p value** | **Spearman's rho** | **p value** | **FDR p value** |
| **PC Score** |  |  |  |  |  |  |  |  |
| *ACTA2* | Pain | Percent change | 0.0333 | 0.9322 | 0.9843 | -0.0333 | 0.9322 | 0.9661 |
|  |  | Delta | 0.0833 | 0.8312 | 0.9529 | -0.0167 | 0.9661 | 0.9661 |
|  | ADL | Percent change | -0.1506 | 0.6989 | 0.9529 | -0.2833 | 0.46 | 0.7763 |
|  |  | Delta | -0.1925 | 0.6198 | 0.9529 | -0.2 | 0.6059 | 0.7790 |
|  | Mean KOOS | Percent change | -0.2667 | 0.4879 | 0.9085 | -0.2833 | 0.46 | 0.7763 |
|  |  | Delta | 0 | 1 | 1.0000 | -0.2176 | 0.5739 | 0.7769 |
| *ANGPT1* | Pain | Percent change | -0.8333 | *0.0053 | 0.2862 | -0.65 | 0.0581 | 0.4636 |
|  |  | Delta | -0.7833 | *0.0125 | 0.3375 | -0.5833 | 0.0992 | 0.4636 |
|  | ADL | Percent change | -0.5523 | 0.1231 | 0.5782 | -0.6833 | *0.0424 | 0.4636 |
|  |  | Delta | -0.6862 | *0.0412 | 0.3708 | -0.7 | *0.0358 | 0.4636 |
|  | Mean KOOS | Percent change | -0.7333 | *0.0246 | 0.3708 | -0.5833 | 0.0992 | 0.4636 |
|  |  | Delta | -0.7167 | *0.0298 | 0.3708 | -0.5941 | 0.0916 | 0.4636 |
| *CCN2* | Pain | Percent change | 0.6167 | 0.0769 | 0.5191 | 0.5667 | 0.1116 | 0.4636 |
|  |  | Delta | 0.7 | *0.0358 | 0.3708 | 0.5167 | 0.1544 | 0.5063 |
|  | ADL | Percent change | 0.5439 | 0.1301 | 0.5782 | 0.5833 | 0.0992 | 0.4636 |
|  |  | Delta | 0.5356 | 0.1373 | 0.5782 | 0.55 | 0.125 | 0.4821 |
|  | Mean KOOS | Percent change | 0.55 | 0.125 | 0.5782 | 0.5667 | 0.1116 | 0.4636 |
|  |  | Delta | 0.6667 | *0.0499 | 0.3849 | 0.5774 | 0.1035 | 0.4636 |
| *CXCL8* | Pain | Percent change | 0.0667 | 0.8647 | 0.9529 | -0.2333 | 0.5457 | 0.7769 |
|  |  | Delta | -0.1167 | 0.765 | 0.9529 | -0.2 | 0.6059 | 0.7790 |
|  | ADL | Percent change | -0.3849 | 0.3063 | 0.8989 | -0.2167 | 0.5755 | 0.7769 |
|  |  | Delta | -0.1423 | 0.715 | 0.9529 | -0.1167 | 0.765 | 0.8606 |
|  | Mean KOOS | Percent change | -0.0667 | 0.8647 | 0.9529 | -0.25 | 0.5165 | 0.7769 |
|  |  | Delta | -0.2833 | 0.46 | 0.9085 | -0.2594 | 0.5003 | 0.7769 |
| *EDN1* | Pain | Percent change | -0.2 | 0.6059 | 0.9529 | -0.5667 | 0.1116 | 0.4636 |
|  |  | Delta | -0.3667 | 0.3317 | 0.8989 | -0.4833 | 0.1875 | 0.5063 |
|  | ADL | Percent change | -0.4268 | 0.252 | 0.8989 | -0.4 | 0.2861 | 0.6437 |
|  |  | Delta | -0.4519 | 0.222 | 0.8563 | -0.4 | 0.2861 | 0.6437 |
|  | Mean KOOS | Percent change | -0.3 | 0.4328 | 0.8989 | -0.5833 | 0.0992 | 0.4636 |
|  |  | Delta | -0.5333 | 0.1392 | 0.5782 | -0.5941 | 0.0916 | 0.4636 |
| *ICAM1* | Pain | Percent change | -0.15 | 0.7001 | 0.9529 | -0.4833 | 0.1875 | 0.5063 |
|  |  | Delta | -0.3333 | 0.3807 | 0.8989 | -0.4167 | 0.2646 | 0.6437 |
|  | ADL | Percent change | -0.1172 | 0.764 | 0.9529 | -0.2333 | 0.5457 | 0.7769 |
|  |  | Delta | -0.3264 | 0.3914 | 0.8989 | -0.2833 | 0.46 | 0.7763 |
|  | Mean KOOS | Percent change | -0.1333 | 0.7324 | 0.9529 | -0.5 | 0.1705 | 0.5063 |
|  |  | Delta | -0.4 | 0.2861 | 0.8989 | -0.5188 | 0.1524 | 0.5063 |
| *PDCD1LG2* | Pain | Percent change | 0.1 | 0.798 | 0.9529 | -0.2167 | 0.5755 | 0.7769 |
|  |  | Delta | -0.1167 | 0.765 | 0.9529 | -0.4167 | 0.2646 | 0.6437 |
|  | ADL | Percent change | -0.1255 | 0.7476 | 0.9529 | -0.0833 | 0.8312 | 0.9160 |
|  |  | Delta | -0.3013 | 0.4308 | 0.8989 | -0.3333 | 0.3807 | 0.7614 |
|  | Mean KOOS | Percent change | -0.0833 | 0.8312 | 0.9529 | -0.1167 | 0.765 | 0.8606 |
|  |  | Delta | -0.1667 | 0.6682 | 0.9529 | -0.1255 | 0.7476 | 0.8606 |
| *THBS1* | Pain | Percent change | -0.0167 | 0.9661 | 0.9843 | -0.3667 | 0.3317 | 0.7165 |
|  |  | Delta | -0.3167 | 0.4064 | 0.8989 | -0.5 | 0.1705 | 0.5063 |
|  | ADL | Percent change | -0.318 | 0.4043 | 0.8989 | -0.2333 | 0.5457 | 0.7769 |
|  |  | Delta | -0.3264 | 0.3914 | 0.8989 | -0.35 | 0.3558 | 0.7390 |
|  | Mean KOOS | Percent change | -0.1333 | 0.7324 | 0.9529 | -0.2833 | 0.46 | 0.7763 |
|  |  | Delta | -0.3833 | 0.3085 | 0.8989 | -0.3096 | 0.4175 | 0.7763 |
| *TNFAIP6* | Pain | Percent change | -0.1 | 0.798 | 0.9529 | 0.0167 | 0.9661 | 0.9661 |
|  |  | Delta | -0.2167 | 0.5755 | 0.9529 | 0.0333 | 0.9322 | 0.9661 |
|  | ADL | Percent change | 0.2762 | 0.472 | 0.9085 | 0.1333 | 0.7324 | 0.8606 |
|  |  | Delta | 0.0502 | 0.8979 | 0.9697 | 0.0333 | 0.9322 | 0.9661 |
|  | Mean KOOS | Percent change | 0.1333 | 0.7324 | 0.9529 | -0.1333 | 0.7324 | 0.8606 |
|  |  | Delta | -0.0167 | 0.9661 | 0.9843 | -0.1172 | 0.764 | 0.8606 |

**Table S3: Discriminant analysis summary for MSC(M)-mediated *in vitro* MΦ polarization.** Samples (count) were assigned to MΦ alone (Solo), Responder, or Non-Responder groups to maximize separation based on multivariate gene expression profiles of each sample. The discriminant analysis reduced all -ddCt values into canonical variables and samples are reclassified into predicted groups based on their gene expression profiles. The fit of the discriminant analysis is described by the entropy R^2^ (values closer to 1 indicate better fit), the fraction of samples that are misclassified into incorrect groups, and by statistical significance testing (Wilks’ Lambda, Pillai’s Trace, Hotelling-Lawley, Roy’s Max Root; *p<0.05).

|  | | | | | **Prob>F** | | | |
| --- | --- | --- | --- | --- | --- | --- | --- | --- |
| **Responder Criteria** | **Count** | **Number misclassified** | **Percent misclassified** | **Entropy R^2^** | **Wilks' Lambda** | **Pillai's Trace** | **Hotelling-Lawley** | **Roy's Max Root** |
| Function-Pain Responder | 28 | 2 | 7.14 | 0.7627 | *0.0002 | *0.0019 | *0.0001 | *<0.0001 |
| Modified Responder classification (patient 10 reclassified) | 28 | 1 | 3.57 | 0.9073 | *<0.0001 | *<0.0001 | *<0.0001 | *<0.0001 |

**Table S4: Significantly differentially expressed microRNAs between MSC(M) derived from Function-Pain Responders versus Non-Responders.** Log2 fold-change values are made relative to Non-Responders (i.e. positive values indicate higher expression in responders).

|  | **baseMean** | **log2FoldChange** | **pvalue** | **padj** |
| --- | --- | --- | --- | --- |
| hsa-miR-642a-5p | 21.1206 | -1.608275508 | 6.31E-07 | 0.000453 |
| hsa-miR-4775 | 65.7219 | 1.055704204 | 8.22E-06 | 0.00295 |
| hsa-miR-210-3p | 91.3405 | -1.253484761 | 1.24E-05 | 0.00296 |
| hsa-miR-338-5p | 10.70128 | 2.161304541 | 8.2E-05 | 0.011781 |
| hsa-miR-3065-5p | 178.3805 | 1.914428105 | 0.000126 | 0.015092 |
| hsa-miR-338-3p | 64.56188 | 2.101813565 | 0.000182 | 0.01634 |
| hsa-miR-19a-3p | 1278.973 | -0.668840623 | 0.00043 | 0.026022 |
| hsa-miR-19b-3p | 4508.064 | -0.657527127 | 0.000435 | 0.026022 |
| hsa-miR-3065-3p | 33.12472 | 2.034926531 | 0.000361 | 0.026022 |
| hsa-miR-302b-3p | 8.832066 | 1.761600415 | 0.000608 | 0.033579 |
| hsa-miR-140-3p | 3219.184 | 0.620396289 | 0.000681 | 0.034929 |
| hsa-miR-31-3p | 143.6965 | -0.787941744 | 0.000924 | 0.044251 |
| hsa-miR-34a-5p | 11171.58 | -0.685564194 | 0.001075 | 0.047701 |
| hsa-miR-483-5p | 16.00379 | 1.631321599 | 0.001129 | 0.047701 |

**Table S5. Correlations between significantly differentially expressed microRNAs in Function-Pain Responder MSC(M) with mRNA levels.** Spearman’s correlation. N=9. Correlations were corrected for multiple comparisons; given exploratory nature of this analysis, correlations that were statistically significant without adjustment for multiple comparisons correction (*p<0.05, FDR>0.1) are marked with an asterisk.

| **microRNA** | **mRNA** | **Spearman ρ** | **Prob>\|ρ\|** | **FDR p value** |
| --- | --- | --- | --- | --- |
| hsa-miR-140-3p | THBS1 | -0.8833 | 0.0016* | 0.1792 |
| hsa-miR-4775 | THBS1 | -0.8117 | 0.0079* | 0.4424 |
| hsa-miR-338-3p | CCN2 | 0.7167 | 0.0298* | 0.6832 |
| hsa-miR-3065-3p | CCN2 | 0.7 | 0.0358* | 0.6832 |
| hsa-miR-140-3p | CXCL8 | -0.7 | 0.0358* | 0.6832 |
| hsa-miR-3065-5p | CCN2 | 0.6778 | 0.0448* | 0.6832 |
| hsa-miR-140-3p | CCN2 | 0.6667 | 0.0499* | 0.6832 |
| hsa-miR-338-3p | CXCL8 | -0.6667 | 0.0499* | 0.6832 |
| hsa-miR-483-5p | THBS1 | -0.65 | 0.0581 | 0.6832 |
| hsa-miR-4775 | CXCL8 | -0.636 | 0.0656 | 0.6832 |
| hsa-miR-210-3p | TNFAIP6 | -0.6333 | 0.0671 | 0.6832 |
| hsa-miR-338-5p | CCN2 | 0.6103 | 0.0809 | 0.7550 |
| hsa-miR-3065-5p | CXCL8 | -0.5858 | 0.0974 | 0.7932 |
| hsa-miR-3065-3p | CXCL8 | -0.5667 | 0.1116 | 0.7932 |
| hsa-miR-3065-3p | PDCD1LG2 | 0.5667 | 0.1116 | 0.7932 |
| hsa-miR-34a-5p | ACTA2 | 0.5333 | 0.1392 | 0.7932 |
| hsa-miR-338-3p | PDCD1LG2 | 0.5333 | 0.1392 | 0.7932 |
| hsa-miR-3065-5p | PDCD1LG2 | 0.5272 | 0.1447 | 0.7932 |
| hsa-miR-338-5p | CXCL8 | -0.5255 | 0.1462 | 0.7932 |
| hsa-miR-4775 | CCN2 | 0.5105 | 0.1603 | 0.7932 |
| hsa-miR-483-5p | PDCD1LG2 | -0.5 | 0.1705 | 0.7932 |
| hsa-miR-4775 | EDN1 | -0.4937 | 0.1768 | 0.7932 |
| hsa-miR-642a-5p | ANGPT1 | 0.4854 | 0.1854 | 0.7932 |
| hsa-miR-4775 | PDCD1LG2 | -0.4854 | 0.1854 | 0.7932 |
| hsa-miR-483-5p | CCN2 | 0.4833 | 0.1875 | 0.7932 |
| hsa-miR-19a-3p | ACTA2 | 0.4667 | 0.2054 | 0.7932 |
| hsa-miR-19b-3p | ACTA2 | 0.4667 | 0.2054 | 0.7932 |
| hsa-miR-483-5p | ANGPT1 | -0.4667 | 0.2054 | 0.7932 |
| hsa-miR-34a-5p | THBS1 | -0.4667 | 0.2054 | 0.7932 |
| hsa-miR-642a-5p | TNFAIP6 | -0.4352 | 0.2418 | 0.8731 |
| hsa-miR-302b-3p | ANGPT1 | -0.41 | 0.273 | 0.8731 |
| hsa-miR-483-5p | EDN1 | -0.4 | 0.2861 | 0.8731 |
| hsa-miR-4775 | TNFAIP6 | 0.3933 | 0.295 | 0.8731 |
| hsa-miR-302b-3p | CCN2 | 0.3849 | 0.3063 | 0.8731 |
| hsa-miR-140-3p | EDN1 | -0.3667 | 0.3317 | 0.8731 |
| hsa-miR-19a-3p | THBS1 | -0.3667 | 0.3317 | 0.8731 |
| hsa-miR-19b-3p | THBS1 | -0.3667 | 0.3317 | 0.8731 |
| hsa-miR-31-3p | TNFAIP6 | -0.3667 | 0.3317 | 0.8731 |
| hsa-miR-3065-5p | ACTA2 | -0.3515 | 0.3537 | 0.8731 |
| hsa-miR-338-5p | PDCD1LG2 | 0.339 | 0.3721 | 0.8731 |
| hsa-miR-31-3p | ACTA2 | 0.3333 | 0.3807 | 0.8731 |
| hsa-miR-210-3p | ANGPT1 | 0.3333 | 0.3807 | 0.8731 |
| hsa-miR-19a-3p | TNFAIP6 | -0.3333 | 0.3807 | 0.8731 |
| hsa-miR-19b-3p | TNFAIP6 | -0.3333 | 0.3807 | 0.8731 |
| hsa-miR-4775 | ANGPT1 | -0.318 | 0.4043 | 0.8731 |
| hsa-miR-3065-3p | ACTA2 | -0.3167 | 0.4064 | 0.8731 |
| hsa-miR-210-3p | PDCD1LG2 | 0.3167 | 0.4064 | 0.8731 |
| hsa-miR-140-3p | PDCD1LG2 | -0.3167 | 0.4064 | 0.8731 |
| hsa-miR-642a-5p | CCN2 | -0.3096 | 0.4175 | 0.8731 |
| hsa-miR-302b-3p | EDN1 | -0.3096 | 0.4175 | 0.8731 |
| hsa-miR-338-5p | ANGPT1 | -0.3051 | 0.4246 | 0.8731 |
| hsa-miR-140-3p | ANGPT1 | -0.3 | 0.4328 | 0.8731 |
| hsa-miR-34a-5p | ANGPT1 | 0.3 | 0.4328 | 0.8731 |
| hsa-miR-34a-5p | PDCD1LG2 | -0.3 | 0.4328 | 0.8731 |
| hsa-miR-338-5p | ACTA2 | -0.2882 | 0.4521 | 0.8731 |
| hsa-miR-338-5p | THBS1 | -0.2882 | 0.4521 | 0.8731 |
| hsa-miR-34a-5p | CXCL8 | -0.2833 | 0.46 | 0.8731 |
| hsa-miR-338-3p | THBS1 | -0.2833 | 0.46 | 0.8731 |
| hsa-miR-302b-3p | CXCL8 | -0.2762 | 0.472 | 0.8731 |
| hsa-miR-3065-3p | ANGPT1 | -0.2667 | 0.4879 | 0.8731 |
| hsa-miR-19a-3p | CXCL8 | -0.2667 | 0.4879 | 0.8731 |
| hsa-miR-19b-3p | CXCL8 | -0.2667 | 0.4879 | 0.8731 |
| hsa-miR-642a-5p | EDN1 | 0.2594 | 0.5003 | 0.8731 |
| hsa-miR-338-3p | ACTA2 | -0.25 | 0.5165 | 0.8731 |
| hsa-miR-338-3p | ANGPT1 | -0.25 | 0.5165 | 0.8731 |
| hsa-miR-3065-5p | ANGPT1 | -0.2427 | 0.5292 | 0.8731 |
| hsa-miR-3065-5p | THBS1 | -0.2427 | 0.5292 | 0.8731 |
| hsa-miR-483-5p | ACTA2 | 0.2333 | 0.5457 | 0.8731 |
| hsa-miR-483-5p | CXCL8 | -0.2333 | 0.5457 | 0.8731 |
| hsa-miR-338-3p | EDN1 | -0.2333 | 0.5457 | 0.8731 |
| hsa-miR-210-3p | EDN1 | 0.2167 | 0.5755 | 0.8952 |
| hsa-miR-3065-3p | THBS1 | -0.2167 | 0.5755 | 0.8952 |
| hsa-miR-302b-3p | ACTA2 | -0.2092 | 0.589 | 0.9036 |
| hsa-miR-338-5p | TNFAIP6 | 0.2034 | 0.5996 | 0.9075 |
| hsa-miR-19a-3p | CCN2 | 0.1833 | 0.6368 | 0.9214 |
| hsa-miR-19b-3p | CCN2 | 0.1833 | 0.6368 | 0.9214 |
| hsa-miR-140-3p | TNFAIP6 | 0.1833 | 0.6368 | 0.9214 |
| hsa-miR-642a-5p | ACTA2 | 0.1674 | 0.6669 | 0.9214 |
| hsa-miR-31-3p | PDCD1LG2 | 0.1667 | 0.6682 | 0.9214 |
| hsa-miR-4775 | ACTA2 | 0.1506 | 0.6989 | 0.9214 |
| hsa-miR-31-3p | CXCL8 | -0.15 | 0.7001 | 0.9214 |
| hsa-miR-31-3p | CCN2 | 0.1333 | 0.7324 | 0.9214 |
| hsa-miR-338-3p | TNFAIP6 | -0.1333 | 0.7324 | 0.9214 |
| hsa-miR-3065-5p | EDN1 | -0.1255 | 0.7476 | 0.9214 |
| hsa-miR-642a-5p | PDCD1LG2 | 0.1255 | 0.7476 | 0.9214 |
| hsa-miR-210-3p | ACTA2 | -0.1167 | 0.765 | 0.9214 |
| hsa-miR-19a-3p | EDN1 | 0.1167 | 0.765 | 0.9214 |
| hsa-miR-19b-3p | EDN1 | 0.1167 | 0.765 | 0.9214 |
| hsa-miR-31-3p | EDN1 | 0.1167 | 0.765 | 0.9214 |
| hsa-miR-642a-5p | CXCL8 | 0.1088 | 0.7806 | 0.9214 |
| hsa-miR-140-3p | ACTA2 | 0.1 | 0.798 | 0.9214 |
| hsa-miR-19a-3p | ANGPT1 | 0.1 | 0.798 | 0.9214 |
| hsa-miR-19b-3p | ANGPT1 | 0.1 | 0.798 | 0.9214 |
| hsa-miR-34a-5p | CCN2 | -0.1 | 0.798 | 0.9214 |
| hsa-miR-3065-3p | EDN1 | -0.1 | 0.798 | 0.9214 |
| hsa-miR-31-3p | THBS1 | -0.1 | 0.798 | 0.9214 |
| hsa-miR-3065-3p | TNFAIP6 | -0.1 | 0.798 | 0.9214 |
| hsa-miR-642a-5p | THBS1 | 0.0837 | 0.8305 | 0.9403 |
| hsa-miR-210-3p | CXCL8 | 0.0833 | 0.8312 | 0.9403 |
| hsa-miR-338-5p | EDN1 | -0.0678 | 0.8624 | 0.9494 |
| hsa-miR-210-3p | CCN2 | -0.0667 | 0.8647 | 0.9494 |
| hsa-miR-210-3p | THBS1 | 0.0667 | 0.8647 | 0.9494 |
| hsa-miR-302b-3p | PDCD1LG2 | 0.0502 | 0.8979 | 0.9582 |
| hsa-miR-3065-5p | TNFAIP6 | -0.0502 | 0.8979 | 0.9582 |
| hsa-miR-34a-5p | EDN1 | -0.05 | 0.8984 | 0.9582 |
| hsa-miR-31-3p | ANGPT1 | 0.0333 | 0.9322 | 0.9849 |
| hsa-miR-302b-3p | TNFAIP6 | 0.0251 | 0.9489 | 0.9926 |
| hsa-miR-19a-3p | PDCD1LG2 | -0.0167 | 0.9661 | 0.9926 |
| hsa-miR-19b-3p | PDCD1LG2 | -0.0167 | 0.9661 | 0.9926 |
| hsa-miR-302b-3p | THBS1 | 0 | 1 | 1 |
| hsa-miR-34a-5p | TNFAIP6 | 0 | 1 | 1 |
| hsa-miR-483-5p | TNFAIP6 | 0 | 1 | 1 |

**Table S6. Correlations between the significantly differentially expressed microRNAs with changes in KOOS (delta and percent change).** N=10 patients. Correlations were corrected for multiple comparisons; given exploratory nature of this analysis, correlations that were statistically significant without multiple comparisons correction (*p<0.05, FDR>0.1) are marked with an asterisk; significant correlations with adjustment for multiple comparisons are bolded (*p<0.05, FDR<0.1). Spearman’s correlation. ADL: function in daily living; KOOS: knee injury and osteoarthritis outcome score; FDR: false discovery rate.

|  | | | **12 month** | | | **24 month** | | |
| --- | --- | --- | --- | --- | --- | --- | --- | --- |
|  | **Follow-up Score** | **Difference Calculation** | **Spearman ρ** | **Prob>\|ρ\|** | **FDR p value** | **Spearman ρ** | **Prob>\|ρ\|** | **FDR p value** |
| hsa-miR-140-3p | ADL | Delta | 0.535 | 0.1111 | 0.2904 | 0.5836 | 0.0765 | 0.2940 |
|  |  | % change | 0.5957 | 0.0692 | 0.2642 | 0.5515 | 0.0984 | 0.2946 |
|  | Mean KOOS | Delta | 0.6242 | 0.0537 | 0.2506 | 0.5829 | 0.077 | 0.2940 |
|  |  | % change | 0.503 | 0.1383 | 0.2904 | 0.5515 | 0.0984 | 0.2946 |
|  | Pain | Delta | 0.5654 | 0.0885 | 0.2791 | 0.6848 | 0.0289* | 0.2114 |
|  |  | % change | 0.4182 | 0.2291 | 0.4095 | 0.6121 | 0.06 | 0.2907 |
| hsa-miR-19a-3p | ADL | Delta | -0.5046 | 0.1369 | 0.2904 | -0.4499 | 0.1921 | 0.3435 |
|  |  | % change | -0.5532 | 0.0972 | 0.2815 | -0.5152 | 0.1276 | 0.3435 |
|  | Mean KOOS | Delta | -0.3939 | 0.26 | 0.4282 | -0.4602 | 0.1808 | 0.3435 |
|  |  | % change | -0.6364 | 0.0479* | 0.2416 | -0.4909 | 0.1497 | 0.3435 |
|  | Pain | Delta | -0.1824 | 0.6141 | 0.6141 | -0.3455 | 0.3282 | 0.4393 |
|  |  | % change | -0.3697 | 0.2931 | 0.4476 | -0.3939 | 0.26 | 0.3766 |
| hsa-miR-19b-3p | ADL | Delta | -0.5046 | 0.1369 | 0.2904 | -0.4499 | 0.1921 | 0.3435 |
|  |  | % change | -0.5532 | 0.0972 | 0.2815 | -0.5152 | 0.1276 | 0.3435 |
|  | Mean KOOS | Delta | -0.3939 | 0.26 | 0.4282 | -0.4602 | 0.1808 | 0.3435 |
|  |  | % change | -0.6364 | 0.0479* | 0.2416 | -0.4909 | 0.1497 | 0.3435 |
|  | Pain | Delta | -0.1824 | 0.6141 | 0.6141 | -0.3455 | 0.3282 | 0.4393 |
|  |  | % change | -0.3697 | 0.2931 | 0.4476 | -0.3939 | 0.26 | 0.3766 |
| hsa-miR-210-3p | ADL | Delta | -0.6565 | 0.0392* | 0.2416 | -0.5957 | 0.0692 | 0.2940 |
|  |  | % change | -0.8632 | 0.0013* | 0.1092 | -0.7333 | 0.0158* | 0.2114 |
|  | Mean KOOS | Delta | -0.6 | 0.0667 | 0.2642 | -0.4724 | 0.168 | 0.3435 |
|  |  | % change | -0.7576 | 0.0111 | 0.1554 | -0.4788 | 0.1615 | 0.3435 |
|  | Pain | Delta | -0.31 | 0.3833 | 0.4897 | -0.4303 | 0.2145 | 0.3465 |
|  |  | % change | -0.503 | 0.1383 | 0.2904 | -0.5515 | 0.0984 | 0.2946 |
| hsa-miR-302b-3p | ADL | Delta | 0.6037 | 0.0646 | 0.2642 | 0.4695 | 0.171 | 0.3435 |
|  |  | % change | 0.8293 | 0.003* | 0.1218 | 0.6505 | 0.0417* | 0.2694 |
|  | Mean KOOS | Delta | 0.5957 | 0.0692 | 0.2642 | 0.4739 | 0.1665 | 0.3435 |
|  |  | % change | 0.6991 | 0.0245* | 0.2159 | 0.5471 | 0.1017 | 0.2946 |
|  | Pain | Delta | 0.2683 | 0.4536 | 0.4964 | 0.2736 | 0.4444 | 0.5258 |
|  |  | % change | 0.4073 | 0.2427 | 0.4247 | 0.4438 | 0.1989 | 0.3435 |
| hsa-miR-3065-3p | ADL | Delta | 0.2128 | 0.5551 | 0.5686 | 0.2371 | 0.5096 | 0.5488 |
|  |  | % change | 0.5106 | 0.1315 | 0.2904 | 0.4303 | 0.2145 | 0.3465 |
|  | Mean KOOS | Delta | 0.3091 | 0.3848 | 0.4897 | 0.3252 | 0.3592 | 0.4503 |
|  |  | % change | 0.3455 | 0.3282 | 0.4753 | 0.3091 | 0.3848 | 0.4685 |
|  | Pain | Delta | 0.2918 | 0.4133 | 0.4964 | 0.1636 | 0.6515 | 0.6515 |
|  |  | % change | 0.2848 | 0.425 | 0.4964 | 0.2242 | 0.5334 | 0.5601 |
| hsa-miR-3065-5p | ADL | Delta | 0.2439 | 0.4971 | 0.5286 | 0.2683 | 0.4536 | 0.5292 |
|  |  | % change | 0.5427 | 0.105 | 0.2845 | 0.462 | 0.1789 | 0.3435 |
|  | Mean KOOS | Delta | 0.3161 | 0.3736 | 0.4897 | 0.3446 | 0.3295 | 0.4393 |
|  |  | % change | 0.3647 | 0.3001 | 0.4502 | 0.3283 | 0.3544 | 0.4503 |
|  | Pain | Delta | 0.2774 | 0.4377 | 0.4964 | 0.1824 | 0.6141 | 0.6291 |
|  |  | % change | 0.2675 | 0.455 | 0.4964 | 0.2371 | 0.5096 | 0.5488 |
| hsa-miR-31-3p | ADL | Delta | -0.4377 | 0.2058 | 0.3929 | -0.4742 | 0.1662 | 0.3435 |
|  |  | % change | -0.462 | 0.1789 | 0.3495 | -0.4424 | 0.2004 | 0.3435 |
|  | Mean KOOS | Delta | -0.3697 | 0.2931 | 0.4476 | -0.497 | 0.1439 | 0.3435 |
|  |  | % change | -0.5273 | 0.1173 | 0.2904 | -0.4788 | 0.1615 | 0.3435 |
|  | Pain | Delta | -0.2979 | 0.4032 | 0.4964 | -0.4909 | 0.1497 | 0.3435 |
|  |  | % change | -0.3455 | 0.3282 | 0.4753 | -0.4182 | 0.2291 | 0.3564 |
| hsa-miR-338-3p | ADL | Delta | 0.231 | 0.5208 | 0.5464 | 0.2492 | 0.4874 | 0.5403 |
|  |  | % change | 0.5228 | 0.121 | 0.2904 | 0.4424 | 0.2004 | 0.3435 |
|  | Mean KOOS | Delta | 0.3333 | 0.3466 | 0.4789 | 0.3497 | 0.3219 | 0.4393 |
|  |  | % change | 0.3333 | 0.3466 | 0.4789 | 0.3333 | 0.3466 | 0.4479 |
|  | Pain | Delta | 0.304 | 0.3932 | 0.4930 | 0.1758 | 0.6272 | 0.6348 |
|  |  | % change | 0.2727 | 0.4458 | 0.4964 | 0.2485 | 0.4888 | 0.5403 |
| hsa-miR-338-5p | ADL | Delta | 0.3262 | 0.3577 | 0.4789 | 0.3508 | 0.3203 | 0.4393 |
|  |  | % change | 0.64 | 0.0462* | 0.2416 | 0.5706 | 0.085 | 0.2946 |
|  | Mean KOOS | Delta | 0.3252 | 0.3592 | 0.4789 | 0.3168 | 0.3725 | 0.4601 |
|  |  | % change | 0.4233 | 0.2228 | 0.4069 | 0.2884 | 0.4191 | 0.5029 |
|  | Pain | Delta | 0.2277 | 0.5269 | 0.5464 | 0.1902 | 0.5987 | 0.6209 |
|  |  | % change | 0.2638 | 0.4614 | 0.4969 | 0.2638 | 0.4614 | 0.5309 |
| hsa-miR-34a-5p | ADL | Delta | -0.3283 | 0.3544 | 0.4789 | -0.3343 | 0.345 | 0.4479 |
|  |  | % change | -0.3951 | 0.2584 | 0.4282 | -0.3939 | 0.26 | 0.3766 |
|  | Mean KOOS | Delta | -0.2727 | 0.4458 | 0.4964 | -0.4111 | 0.2379 | 0.3633 |
|  |  | % change | -0.4909 | 0.1497 | 0.2994 | -0.4182 | 0.2291 | 0.3564 |
|  | Pain | Delta | -0.2736 | 0.4444 | 0.4964 | -0.2242 | 0.5334 | 0.5601 |
|  |  | % change | -0.3818 | 0.2763 | 0.4463 | -0.2485 | 0.4888 | 0.5403 |
| hsa-miR-4775 | ADL | Delta | 0.6951 | 0.0257* | 0.2159 | 0.6982 | 0.0247* | 0.2114 |
|  |  | % change | 0.7683 | 0.0094* | 0.1554 | 0.7052 | 0.0227* | 0.2114 |
|  | Mean KOOS | Delta | 0.7295 | 0.0166* | 0.1743 | 0.6277 | 0.052 | 0.2907 |
|  |  | % change | 0.6626 | 0.0368* | 0.2416 | 0.6079 | 0.0623 | 0.2907 |
|  | Pain | Delta | 0.5427 | 0.105 | 0.2845 | 0.7234 | 0.018* | 0.2114 |
|  |  | % change | 0.4924 | 0.1482 | 0.2994 | 0.7173 | 0.0195 | 0.2114 |
| hsa-miR-483-5p | ADL | Delta | 0.5714 | 0.0844 | 0.2791 | 0.6809 | 0.0302* | 0.2114 |
|  |  | % change | 0.2675 | 0.455 | 0.4964 | 0.4303 | 0.2145 | 0.3465 |
|  | Mean KOOS | Delta | 0.6364 | 0.0479* | 0.2416 | 0.681 | 0.0302* | 0.2114 |
|  |  | % change | 0.4303 | 0.2145 | 0.4004 | 0.6121 | 0.06 | 0.2907 |
|  | Pain | Delta | 0.7964 | 0.0058* | 0.1218 | **0.8788** | **0.0008*** | **0.0672** |
|  |  | % change | 0.5636 | 0.0897 | 0.2791 | 0.7333 | 0.0158* | 0.2114 |
| hsa-miR-642a-5p | ADL | Delta | -0.6341 | 0.0489 | 0.2416 | -0.686 | 0.0285* | 0.2114 |
|  |  | % change | -0.7988 | 0.0056* | 0.1218 | -0.7842 | 0.0072* | 0.2114 |
|  | Mean KOOS | Delta | -0.5775 | 0.0804 | 0.2791 | -0.6216 | 0.0551 | 0.2907 |
|  |  | % change | -0.7295 | 0.0166* | 0.1743 | -0.5654 | 0.0885 | 0.2946 |
|  | Pain | Delta | -0.5061 | 0.1356 | 0.2904 | -0.5471 | 0.1017 | 0.2946 |
|  |  | % change | -0.5836 | 0.0765 | 0.2791 | -0.5897 | 0.0728 | 0.2940 |

**Table S7. Correlations between baseline KOOS and changes in KOOS (percent change or delta values) at 12- and 24-months.** N=12 patients. Correlations were corrected for multiple comparisons; given exploratory nature of this analysis, correlations that were statistically significant without multiple comparisons correction (*p<0.05, FDR>0.1) are marked with an asterisk. Spearman’s correlation. ADL: function in daily living; Sports/Rec: function in sport and recreation; QOL: knee-related quality of life; KOOS: knee injury and osteoarthritis outcome score.

|  | | | **12 Months** | | | **24 Months** | | |
| --- | --- | --- | --- | --- | --- | --- | --- | --- |
| **Baseline Score** | **Follow-up Score** | **Difference calculation** | **Spearman's rho** | **p value** | **FDR p value** | **Spearman's rho** | **p value** | **FDR p value** |
| Symptom | Pain | Percent change | 0.0839 | 0.7954 | 0.8629 | 0.3427 | 0.2756 | 0.3969 |
|  |  | Delta | 0.3328 | 0.2906 | 0.6030 | 0.5219 | 0.0818 | 0.3969 |
|  | ADL | Percent change | 0.2487 | 0.4357 | 0.6775 | 0.2238 | 0.4845 | 0.5814 |
|  |  | Delta | 0.3117 | 0.3239 | 0.6137 | 0.4063 | 0.19 | 0.3969 |
|  | Mean KOOS | Percent change | 0.0909 | 0.7787 | 0.8629 | 0.3287 | 0.2969 | 0.4111 |
|  |  | Delta | 0.3287 | 0.2969 | 0.6030 | 0.4085 | 0.1874 | 0.3969 |
| Pain | Pain | Percent change | 0.0736 | 0.8203 | 0.8629 | 0.1681 | 0.6015 | 0.6538 |
|  |  | Delta | 0.4053 | 0.1912 | 0.6030 | 0.4684 | 0.1246 | 0.3969 |
|  | ADL | Percent change | -0.0561 | 0.8624 | 0.8629 | -0.0525 | 0.8712 | 0.8712 |
|  |  | Delta | 0.1123 | 0.7283 | 0.8629 | 0.1632 | 0.6124 | 0.6538 |
|  | Mean KOOS | Percent change | -0.0876 | 0.7867 | 0.8629 | 0.1296 | 0.6881 | 0.7078 |
|  |  | Delta | 0.1191 | 0.7124 | 0.8629 | 0.2328 | 0.4665 | 0.5814 |
| ADL | Pain | Percent change | 0.1608 | 0.6175 | 0.8550 | 0.2028 | 0.5273 | 0.6123 |
|  |  | Delta | 0.3573 | 0.2542 | 0.6030 | 0.3538 | 0.2593 | 0.3969 |
|  | ADL | Percent change | 0.2417 | 0.4492 | 0.6775 | 0.3497 | 0.2652 | 0.3969 |
|  |  | Delta | 0.4974 | 0.0999 | 0.6030 | 0.4939 | 0.1027 | 0.3969 |
|  | Mean KOOS | Percent change | 0.2308 | 0.4705 | 0.6775 | 0.3776 | 0.2262 | 0.3969 |
|  |  | Delta | 0.3497 | 0.2652 | 0.6030 | 0.4296 | 0.1634 | 0.3969 |
| Sports/Rec | Pain | Percent change | 0.3937 | 0.2055 | 0.6030 | 0.3691 | 0.2377 | 0.3969 |
|  |  | Delta | 0.5599 | 0.0584 | 0.6030 | 0.4877 | 0.1078 | 0.3969 |
|  | ADL | Percent change | 0.3398 | 0.2799 | 0.6030 | 0.4605 | 0.132 | 0.3969 |
|  |  | Delta | 0.6056 | *0.0369 | 0.6030 | 0.5845 | *0.0459 | 0.3969 |
|  | Mean KOOS | Percent change | 0.3796 | 0.2236 | 0.6030 | 0.5448 | 0.067 | 0.3969 |
|  |  | Delta | 0.5202 | 0.0829 | 0.6030 | 0.6018 | *0.0384 | 0.3969 |
| QOL | Pain | Percent change | 0.2963 | 0.3497 | 0.6295 | 0.4445 | 0.1477 | 0.3969 |
|  |  | Delta | 0.4806 | 0.1138 | 0.6030 | 0.5477 | 0.0653 | 0.3969 |
|  | ADL | Percent change | 0.4452 | 0.1469 | 0.6030 | 0.3915 | 0.2081 | 0.3969 |
|  |  | Delta | 0.3445 | 0.2728 | 0.6030 | 0.3569 | 0.2548 | 0.3969 |
|  | Mean KOOS | Percent change | 0.2857 | 0.368 | 0.6309 | 0.4339 | 0.1588 | 0.3969 |
|  |  | Delta | 0.4692 | 0.1239 | 0.6030 | 0.5044 | 0.0944 | 0.3969 |
| Mean KOOS | Pain | Percent change | 0.0559 | 0.8629 | 0.8629 | 0.1608 | 0.6175 | 0.6538 |
|  |  | Delta | 0.3257 | 0.3015 | 0.6030 | 0.3783 | 0.2253 | 0.3969 |
|  | ADL | Percent change | 0.1366 | 0.6721 | 0.8629 | 0.2238 | 0.4845 | 0.5814 |
|  |  | Delta | 0.3643 | 0.2444 | 0.6030 | 0.4028 | 0.1942 | 0.3969 |
|  | Mean KOOS | Percent change | 0.0769 | 0.8122 | 0.8629 | 0.2867 | 0.3663 | 0.4884 |
|  |  | Delta | 0.2378 | 0.4568 | 0.6775 | 0.3662 | 0.2417 | 0.3969 |

**Table S8: Correlations between baseline MARS physical activity scores and changes in KOOS (percent change or delta values).** N=12 patients. Significant correlations with adjustments for multiple comparisons (*p<0.05; FDR<0.1) are bolded. Spearman’s correlation. ADL: function in daily living; Sports/Rec: function in sport and recreation; QOL: knee-related quality of life; KOOS: knee injury and osteoarthritis outcome score; MARS; Marx Activity Rating Scale.

|  | | **12 Month** | | | **24 Month** | | |
| --- | --- | --- | --- | --- | --- | --- | --- |
| **Follow-up Score** | **Difference calculation** | **Spearman's rho** | **p value** | **FDR p value** | **Spearman's rho** | **p value** | **FDR p value** |
| Pain | Percent change | **-0.7502** | ***0.0049** | **0.0147** | 0.207 | 0.5186 | 0.8843 |
|  | Delta | **-0.7777** | ***0.0029** | **0.0147** | 0.0984 | 0.7609 | 0.8843 |
| ADL | Percent change | 0.2882 | 0.3636 | 0.43632 | 0.1298 | 0.6876 | 0.8843 |
|  | Delta | 0.225 | 0.4821 | 0.4821 | 0.1617 | 0.6156 | 0.8843 |
| QOL | Percent change | -0.3402 | 0.2792 | 0.4188 | 0.1742 | 0.5883 | 0.8843 |
|  | Delta | -0.4449 | 0.1472 | 0.2944 | 0.0472 | 0.8843 | 0.8843 |

**Table S9. Correlations between baseline MRI WORMS and changes in KOOS (percent change or delta values).** N=12 patients. Spearman’s correlation. ADL: function in daily living; KOOS: knee injury and osteoarthritis outcome score.

|  | | **12 Month** | | | **24 Month** | | |
| --- | --- | --- | --- | --- | --- | --- | --- |
| **Follow-up Score** | **Difference calculation** | **Spearman's rho** | **p value** | **FDR p value** | **Spearman's rho** | **p value** | **FDR p value** |
| Pain | Delta | -0.1541 | 0.6325 | 0.8861 | 0.014 | 0.9655 | 0.9655 |
|  | Percent change | -0.2308 | 0.4705 | 0.8861 | -0.0699 | 0.829 | 0.9044 |
| ADL | Delta | -0.2732 | 0.3902 | 0.8861 | -0.1716 | 0.5938 | 0.8861 |
|  | Percent change | -0.1821 | 0.571 | 0.8861 | -0.2797 | 0.3786 | 0.8861 |
| Mean KOOS | Delta | -0.1888 | 0.5567 | 0.8861 | -0.0916 | 0.7772 | 0.9044 |
|  | Percent change | -0.3077 | 0.3306 | 0.8861 | -0.1399 | 0.6646 | 0.8861 |

**Table S10. Correlations between baseline MRI synovitis scores and changes in KOOS (percent change or delta values).** N=10 patients. Spearman’s correlation. ADL: function in daily living; KOOS: knee injury and osteoarthritis outcome score.

|  |  | **12 Month** | | | **24 Month** | | |
| --- | --- | --- | --- | --- | --- | --- | --- |
| **Follow-up Score** | **Difference calculation** | **Spearman's rho** | **p value** | **FDR p value** | **Spearman's rho** | **p value** | **FDR p value** |
| Pain | Delta | -0.0092 | 0.9799 | 0.9799 | -0.4557 | 0.1857 | 0.3279 |
|  | Percent change | -0.1829 | 0.613 | 0.6687 | -0.4329 | 0.2114 | 0.3279 |
| ADL | Delta | -0.3089 | 0.3852 | 0.5136 | -0.4329 | 0.2114 | 0.3279 |
|  | Percent change | -0.4954 | 0.1454 | 0.3279 | -0.4756 | 0.1647 | 0.3279 |
| Mean KOOS | Delta | -0.2805 | 0.4325 | 0.5190 | -0.5107 | 0.1314 | 0.3279 |
|  | Percent change | -0.4817 | 0.1586 | 0.3279 | -0.4268 | 0.2186 | 0.3279 |

**Table S11: Correlations between changes in KOOS (percent change or delta values) with baseline levels of local synovial fluid metabolic, angiogenic, and extracellular matrix-associated biomarkers.** N=8-9 patients. Correlations were corrected for multiple comparisons; given exploratory nature of this analysis, correlations that were statistically significant without multiple comparisons correction (*p<0.05, FDR>0.1) are marked with an asterisk. Spearman’s correlation. ADL: function in daily living; KOOS: knee injury and osteoarthritis outcome score; HGF: hepatocyte growth factor; MMP: matrix metalloproteinase; TIMP1: tissue inhibitor of metalloproteinases 1; VEGFA: vascular endothelial growth factor A.

|  | | | **6 Month** | | | **12 Month** | | | **24 Month** | | |
| --- | --- | --- | --- | --- | --- | --- | --- | --- | --- | --- | --- |
| **Biomarker** | **Follow-up Score** | **Difference calculation** | **Spearman's rho** | **p value** | **FDR p value** | **Spearman's rho** | **p value** | **FDR p value** | **Spearman's rho** | **p value** | **FDR p value** |
| Adiponectin | Pain | Percent change | 0.1 | 0.798 | 0.9857 | 0.0167 | 0.9661 | 0.9810 | 0.2167 | 0.5755 | 1.0000 |
|  |  | Delta | 0.2092 | 0.589 | 0.9857 | 0.1 | 0.798 | 0.9810 | 0.3333 | 0.3807 | 1.0000 |
|  | ADL | Percent change | 0.0667 | 0.8647 | 0.9857 | 0.0833 | 0.8312 | 0.9810 | -0.0833 | 0.8312 | 1.0000 |
|  |  | Delta | 0.0921 | 0.8138 | 0.9857 | 0.0667 | 0.8647 | 0.9810 | 0.0833 | 0.8312 | 1.0000 |
|  | Mean KOOS | Percent change | 0.2 | 0.6059 | 0.9857 | 0.2333 | 0.5457 | 0.9810 | 0.1833 | 0.6368 | 1.0000 |
|  |  | Delta | 0.0833 | 0.8312 | 0.9857 | 0.0333 | 0.9322 | 0.9810 | 0.2667 | 0.4879 | 1.0000 |
| Adipsin | Pain | Percent change | 0.25 | 0.5165 | 0.9857 | 0.0833 | 0.8312 | 0.9810 | 0.2 | 0.6059 | 1.0000 |
|  |  | Delta | 0.1172 | 0.764 | 0.9857 | -0.2333 | 0.5457 | 0.9810 | 0.0167 | 0.9661 | 1.0000 |
|  | ADL | Percent change | 0.4667 | 0.2054 | 0.9857 | 0.4 | 0.2861 | 0.9810 | 0.4 | 0.2861 | 1.0000 |
|  |  | Delta | 0.1339 | 0.7313 | 0.9857 | 0.0167 | 0.9661 | 0.9810 | 0.1833 | 0.6368 | 1.0000 |
|  | Mean KOOS | Percent change | 0.4 | 0.2861 | 0.9857 | 0.4 | 0.2861 | 0.9810 | 0.15 | 0.7001 | 1.0000 |
|  |  | Delta | 0.1833 | 0.6368 | 0.9857 | 0.1167 | 0.765 | 0.9810 | 0.1167 | 0.765 | 1.0000 |
| Leptin | Pain | Percent change | 0.05 | 0.8984 | 0.9857 | 0.0667 | 0.8647 | 0.9810 | -0.3833 | 0.3085 | 1.0000 |
|  |  | Delta | 0.1255 | 0.7476 | 0.9857 | 0.2333 | 0.5457 | 0.9810 | -0.2167 | 0.5755 | 1.0000 |
|  | ADL | Percent change | -0.3667 | 0.3317 | 0.9857 | -0.1833 | 0.6368 | 0.9810 | -0.35 | 0.3558 | 1.0000 |
|  |  | Delta | -0.1925 | 0.6198 | 0.9857 | 0.0167 | 0.9661 | 0.9810 | -0.1667 | 0.6682 | 1.0000 |
|  | Mean KOOS | Percent change | -0.2 | 0.6059 | 0.9857 | -0.2 | 0.6059 | 0.9810 | -0.1333 | 0.7324 | 1.0000 |
|  |  | Delta | -0.1667 | 0.6682 | 0.9857 | 0.0167 | 0.9661 | 0.9810 | -0.1833 | 0.6368 | 1.0000 |
| Resistin | Pain | Percent change | -0.25 | 0.5165 | 0.9857 | -0.15 | 0.7001 | 0.9810 | 0.25 | 0.5165 | 1.0000 |
|  |  | Delta | -0.0418 | 0.9149 | 0.9857 | 0.1333 | 0.7324 | 0.9810 | 0.3833 | 0.3085 | 1.0000 |
|  | ADL | Percent change | 0.1667 | 0.6682 | 0.9857 | 0.1167 | 0.765 | 0.9810 | 0.2167 | 0.5755 | 1.0000 |
|  |  | Delta | 0.4603 | 0.2125 | 0.9857 | 0.35 | 0.3558 | 0.9810 | 0.3 | 0.4328 | 1.0000 |
|  | Mean KOOS | Percent change | 0.1 | 0.798 | 0.9857 | 0.1333 | 0.7324 | 0.9810 | 0.2667 | 0.4879 | 1.0000 |
|  |  | Delta | 0.3 | 0.4328 | 0.9857 | 0.1167 | 0.765 | 0.9810 | 0.3833 | 0.3085 | 1.0000 |
| HGF | Pain | Percent change | -0.4524 | 0.2604 | 0.9857 | -0.7143 | *0.0465 | 0.9570 | -0.381 | 0.3518 | 1.0000 |
|  |  | Delta | -0.5714 | 0.139 | 0.9857 | -0.7857 | *0.0208 | 0.6864 | -0.5476 | 0.16 | 1.0000 |
|  | ADL | Percent change | -0.0476 | 0.9108 | 0.9857 | -0.4048 | 0.3199 | 0.9810 | -0.381 | 0.3518 | 1.0000 |
|  |  | Delta | -0.2156 | 0.6081 | 0.9857 | -0.5238 | 0.1827 | 0.9810 | -0.381 | 0.3518 | 1.0000 |
|  | Mean KOOS | Percent change | -0.3333 | 0.4198 | 0.9857 | -0.5476 | 0.16 | 0.9810 | -0.4762 | 0.2329 | 1.0000 |
|  |  | Delta | -0.381 | 0.3518 | 0.9857 | -0.5952 | 0.1195 | 0.9810 | -0.4286 | 0.2894 | 1.0000 |
| VEGFA | Pain | Percent change | -0.4762 | 0.2329 | 0.9857 | -0.8095 | *0.0149 | 0.6864 | -0.4048 | 0.3199 | 1.0000 |
|  |  | Delta | -0.619 | 0.1017 | 0.9857 | -0.6905 | 0.058 | 0.9570 | -0.4048 | 0.3199 | 1.0000 |
|  | ADL | Percent change | 0.0238 | 0.9554 | 0.9857 | -0.2143 | 0.6103 | 0.9810 | -0.4524 | 0.2604 | 1.0000 |
|  |  | Delta | -0.1796 | 0.6703 | 0.9857 | -0.5238 | 0.1827 | 0.9810 | -0.381 | 0.3518 | 1.0000 |
|  | Mean KOOS | Percent change | -0.381 | 0.3518 | 0.9857 | -0.5714 | 0.139 | 0.9810 | -0.4762 | 0.2329 | 1.0000 |
|  |  | Delta | -0.381 | 0.3518 | 0.9857 | -0.5238 | 0.1827 | 0.9810 | -0.4048 | 0.3199 | 1.0000 |
| MMP13 | Pain | Percent change | 0 | 1 | 1.0000 | -0.0238 | 0.9554 | 0.9810 | -0.0714 | 0.8665 | 1.0000 |
|  |  | Delta | 0.1429 | 0.7358 | 0.9857 | -0.0238 | 0.9554 | 0.9810 | 0.0476 | 0.9108 | 1.0000 |
|  | ADL | Percent change | -0.381 | 0.3518 | 0.9857 | -0.2619 | 0.5309 | 0.9810 | -0.2619 | 0.5309 | 1.0000 |
|  |  | Delta | -0.1198 | 0.7776 | 0.9857 | -0.0714 | 0.8665 | 0.9810 | -0.0952 | 0.8225 | 1.0000 |
|  | Mean KOOS | Percent change | 0.0238 | 0.9554 | 0.9857 | 0.0952 | 0.8225 | 0.9810 | 0 | 1 | 1.0000 |
|  |  | Delta | -0.0952 | 0.8225 | 0.9857 | -0.1905 | 0.6514 | 0.9810 | 0.0476 | 0.9108 | 1.0000 |
| MMP13 | Pain | Percent change | -0.0935 | 0.8257 | 0.9857 | -0.3429 | 0.4057 | 0.9810 | 0.2338 | 0.5773 | 1.0000 |
|  |  | Delta | -0.2338 | 0.5773 | 0.9857 | -0.2338 | 0.5773 | 0.9810 | 0.2338 | 0.5773 | 1.0000 |
|  | ADL | Percent change | 0.5455 | 0.1619 | 0.9857 | 0.3273 | 0.4287 | 0.9810 | 0.0156 | 0.9708 | 1.0000 |
|  |  | Delta | 0.149 | 0.7248 | 0.9857 | -0.265 | 0.5259 | 0.9810 | -0.0156 | 0.9708 | 1.0000 |
|  | Mean KOOS | Percent change | 0.0156 | 0.9708 | 0.9857 | -0.0935 | 0.8257 | 0.9810 | -0.0156 | 0.9708 | 1.0000 |
|  |  | Delta | -0.0156 | 0.9708 | 0.9857 | -0.0935 | 0.8257 | 0.9810 | 0.0935 | 0.8257 | 1.0000 |
| MMP3 | Pain | Percent change | -0.119 | 0.7789 | 0.9857 | -0.1429 | 0.7358 | 0.9810 | -0.0476 | 0.9108 | 1.0000 |
|  |  | Delta | 0.0238 | 0.9554 | 0.9857 | -0.1429 | 0.7358 | 0.9810 | 0.0714 | 0.8665 | 1.0000 |
|  | ADL | Percent change | -0.3333 | 0.4198 | 0.9857 | -0.2857 | 0.4927 | 0.9810 | -0.3571 | 0.3851 | 1.0000 |
|  |  | Delta | -0.1916 | 0.6494 | 0.9857 | -0.2381 | 0.5702 | 0.9810 | -0.2143 | 0.6103 | 1.0000 |
|  | Mean KOOS | Percent change | -0.0714 | 0.8665 | 0.9857 | 0 | 1 | 1.0000 | -0.0952 | 0.8225 | 1.0000 |
|  |  | Delta | -0.2143 | 0.6103 | 0.9857 | -0.3333 | 0.4198 | 0.9810 | 0 | 1 | 1.0000 |
| MMP9 | Pain | Percent change | 0.1952 | 0.6432 | 0.9857 | -0.1708 | 0.686 | 0.9810 | 0.2196 | 0.6013 | 1.0000 |
|  |  | Delta | 0.0244 | 0.9543 | 0.9857 | -0.2684 | 0.5204 | 0.9810 | 0.0244 | 0.9543 | 1.0000 |
|  | ADL | Percent change | 0.5855 | 0.1272 | 0.9857 | 0.2684 | 0.5204 | 0.9810 | 0.244 | 0.5604 | 1.0000 |
|  |  | Delta | 0.3313 | 0.4227 | 0.9857 | -0.0244 | 0.9543 | 0.9810 | 0.2196 | 0.6013 | 1.0000 |
|  | Mean KOOS | Percent change | 0.3416 | 0.4076 | 0.9857 | 0.122 | 0.7735 | 0.9810 | 0.1464 | 0.7294 | 1.0000 |
|  |  | Delta | 0.2196 | 0.6013 | 0.9857 | 0.0244 | 0.9543 | 0.9810 | 0.1708 | 0.686 | 1.0000 |
| TIMP1 | Pain | Percent change | -0.1905 | 0.6514 | 0.9857 | -0.4762 | 0.2329 | 0.9810 | -0.0476 | 0.9108 | 1.0000 |
|  |  | Delta | -0.2619 | 0.5309 | 0.9857 | -0.4524 | 0.2604 | 0.9810 | -0.0476 | 0.9108 | 1.0000 |
|  | ADL | Percent change | 0.1429 | 0.7358 | 0.9857 | -0.0238 | 0.9554 | 0.9810 | -0.2619 | 0.5309 | 1.0000 |
|  |  | Delta | -0.0958 | 0.8215 | 0.9857 | -0.4286 | 0.2894 | 0.9810 | -0.2143 | 0.6103 | 1.0000 |
|  | Mean KOOS | Percent change | -0.0714 | 0.8665 | 0.9857 | -0.1667 | 0.6932 | 0.9810 | -0.1905 | 0.6514 | 1.0000 |
|  |  | Delta | -0.2143 | 0.6103 | 0.9857 | -0.3571 | 0.3851 | 0.9810 | -0.0952 | 0.8225 | 1.0000 |

**Table S12: Correlations between changes in KOOS (percent change or delta values) with baseline levels of local synovial fluid inflammatory biomarkers.** N=8 patients. No significant correlations. Spearman’s correlation. ADL: function in daily living; KOOS: knee injury and osteoarthritis outcome score. CCL2: chemokine ligand 2; CX3CL1: chemokine (C-X3-C motif) ligand 1; CXCL1: chemokine (C-X-C motif) ligand 1; IL12p40: IL: interleukin; sCD14: soluble CD14; sCD163: soluble CD163.

|  | | | **6 Month** | | | **12 Month** | | | **24 Month** | | |
| --- | --- | --- | --- | --- | --- | --- | --- | --- | --- | --- | --- |
| **Biomarker** | **Follow-up Score** | **Difference calculation** | **Spearman's rho** | **p value** | **FDR p value** | **Spearman's rho** | **p value** | **FDR p value** | **Spearman's rho** | **p value** | **FDR p value** |
| CCL2 | Pain | Percent change | -0.1667 | 0.6932 | 0.9715 | -0.2381 | 0.5702 | 0.9757 | -0.2857 | 0.4927 | 0.9983 |
|  |  | Delta | -0.1429 | 0.7358 | 0.9715 | -0.2143 | 0.6103 | 0.9757 | -0.1429 | 0.7358 | 0.9983 |
|  | ADL | Percent change | -0.1905 | 0.6514 | 0.9715 | -0.0238 | 0.9554 | 0.9757 | -0.4048 | 0.3199 | 0.9983 |
|  |  | Delta | -0.4192 | 0.3013 | 0.9715 | -0.4048 | 0.3199 | 0.9757 | -0.381 | 0.3518 | 0.9983 |
|  | Mean KOOS | Percent change | -0.1905 | 0.6514 | 0.9715 | -0.0952 | 0.8225 | 0.9757 | -0.3095 | 0.4556 | 0.9983 |
|  |  | Delta | -0.381 | 0.3518 | 0.9715 | -0.2857 | 0.4927 | 0.9757 | -0.2857 | 0.4927 | 0.9983 |
| CX3CL1 | Pain | Percent change | -0.2474 | 0.5546 | 0.9715 | -0.4124 | 0.31 | 0.9757 | 0.2474 | 0.5546 | 0.9983 |
|  |  | Delta | -0.2474 | 0.5546 | 0.9715 | -0.2474 | 0.5546 | 0.9757 | 0.2474 | 0.5546 | 0.9983 |
|  | ADL | Percent change | 0.0825 | 0.8461 | 0.9715 | -0.2474 | 0.5546 | 0.9757 | -0.0825 | 0.8461 | 0.9983 |
|  |  | Delta | 0.3319 | 0.4219 | 0.9715 | -0.0825 | 0.8461 | 0.9757 | 0.0825 | 0.8461 | 0.9983 |
|  | Mean KOOS | Percent change | -0.0825 | 0.8461 | 0.9715 | -0.2474 | 0.5546 | 0.9757 | 0.0825 | 0.8461 | 0.9983 |
|  |  | Delta | 0.0825 | 0.8461 | 0.9715 | -0.2474 | 0.5546 | 0.9757 | 0.2474 | 0.5546 | 0.9983 |
| CXCL1 | Pain | Percent change | -0.2474 | 0.5546 | 0.9715 | -0.4124 | 0.31 | 0.9757 | 0.2474 | 0.5546 | 0.9983 |
|  |  | Delta | -0.2474 | 0.5546 | 0.9715 | -0.2474 | 0.5546 | 0.9757 | 0.2474 | 0.5546 | 0.9983 |
|  | ADL | Percent change | 0.0825 | 0.8461 | 0.9715 | -0.2474 | 0.5546 | 0.9757 | -0.0825 | 0.8461 | 0.9983 |
|  |  | Delta | 0.3319 | 0.4219 | 0.9715 | -0.0825 | 0.8461 | 0.9757 | 0.0825 | 0.8461 | 0.9983 |
|  | Mean KOOS | Percent change | -0.0825 | 0.8461 | 0.9715 | -0.2474 | 0.5546 | 0.9757 | 0.0825 | 0.8461 | 0.9983 |
|  |  | Delta | 0.0825 | 0.8461 | 0.9715 | -0.2474 | 0.5546 | 0.9757 | 0.2474 | 0.5546 | 0.9983 |
| IL12p40 | Pain | Percent change | 0.1317 | 0.7558 | 0.9715 | -0.2275 | 0.5878 | 0.9757 | -0.1078 | 0.7995 | 0.9983 |
|  |  | Delta | 0.0479 | 0.9103 | 0.9715 | -0.0838 | 0.8435 | 0.9757 | -0.012 | 0.9775 | 0.9983 |
|  | ADL | Percent change | 0.2635 | 0.5284 | 0.9715 | 0.2036 | 0.6287 | 0.9757 | -0.0479 | 0.9103 | 0.9983 |
|  |  | Delta | 0.2892 | 0.4873 | 0.9715 | 0.1198 | 0.7776 | 0.9757 | 0.1437 | 0.7342 | 0.9983 |
|  | Mean KOOS | Percent change | 0.1796 | 0.6703 | 0.9715 | 0.024 | 0.9551 | 0.9757 | 0.0359 | 0.9327 | 0.9983 |
|  |  | Delta | 0.1437 | 0.7342 | 0.9715 | 0.024 | 0.9551 | 0.9757 | 0.0599 | 0.888 | 0.9983 |
| IL6 | Pain | Percent change | -0.119 | 0.7789 | 1.0000 | -0.1905 | 0.6514 | 0.9757 | 0.0476 | 0.9108 | 0.9983 |
|  |  | Delta | 0 | 1 | 0.9715 | -0.0952 | 0.8225 | 0.9757 | 0.1905 | 0.6514 | 0.9983 |
|  | ADL | Percent change | -0.2381 | 0.5702 | 0.9983 | -0.2381 | 0.5702 | 0.9757 | -0.2857 | 0.4927 | 0.9983 |
|  |  | Delta | -0.012 | 0.9775 | 0.9715 | -0.1429 | 0.7358 | 0.9757 | -0.0952 | 0.8225 | 0.9983 |
|  | Mean KOOS | Percent change | -0.0476 | 0.9108 | 0.9715 | -0.0238 | 0.9554 | 0.9757 | 0 | 1 | 1.0000 |
|  |  | Delta | -0.0952 | 0.8225 | 0.9966 | -0.2619 | 0.5309 | 0.9757 | 0.119 | 0.7789 | 0.9983 |
| IL8 | Pain | Percent change | 0.024 | 0.9551 | 0.9715 | -0.3114 | 0.4528 | 0.9757 | 0.0719 | 0.8657 | 0.9983 |
|  |  | Delta | -0.1317 | 0.7558 | 0.9715 | -0.4551 | 0.2572 | 0.9757 | -0.1677 | 0.6915 | 0.9983 |
|  | ADL | Percent change | 0.3832 | 0.3487 | 0.9715 | 0.024 | 0.9551 | 0.9757 | 0.0958 | 0.8215 | 0.9983 |
|  |  | Delta | 0.1687 | 0.6897 | 0.9715 | -0.1796 | 0.6703 | 0.9757 | 0.0599 | 0.888 | 0.9983 |
|  | Mean KOOS | Percent change | 0.1796 | 0.6703 | 0.9715 | -0.0479 | 0.9103 | 0.9757 | -0.012 | 0.9775 | 0.9983 |
|  |  | Delta | 0.0599 | 0.888 | 0.9715 | -0.1677 | 0.6915 | 0.9757 | 0.012 | 0.9775 | 0.9983 |
| sCD14 | Pain | Percent change | -0.0714 | 0.8665 | 0.9715 | 0.0476 | 0.9108 | 0.9757 | -0.3333 | 0.4198 | 0.9983 |
|  |  | Delta | 0.0476 | 0.9108 | 0.9715 | 0.119 | 0.7789 | 0.9757 | -0.0952 | 0.8225 | 0.9983 |
|  | ADL | Percent change | -0.4286 | 0.2894 | 0.9715 | -0.0714 | 0.8665 | 0.9757 | -0.3571 | 0.3851 | 0.9983 |
|  |  | Delta | -0.3952 | 0.3325 | 0.9715 | -0.119 | 0.7789 | 0.9757 | -0.2857 | 0.4927 | 0.9983 |
|  | Mean KOOS | Percent change | -0.1905 | 0.6514 | 0.9715 | 0 | 1 | 1.0000 | -0.2143 | 0.6103 | 0.9983 |
|  |  | Delta | -0.2857 | 0.4927 | 0.9715 | -0.0952 | 0.8225 | 0.9757 | -0.2381 | 0.5702 | 0.9983 |
| sCD163 | Pain | Percent change | 0.619 | 0.1017 | 0.9715 | 0.5 | 0.207 | 0.9757 | 0.2857 | 0.4927 | 0.9983 |
|  |  | Delta | 0.6667 | 0.071 | 0.9715 | 0.381 | 0.3518 | 0.9757 | 0.2381 | 0.5702 | 0.9983 |
|  | ADL | Percent change | 0.2143 | 0.6103 | 0.9715 | 0.2381 | 0.5702 | 0.9757 | 0.4048 | 0.3199 | 0.9983 |
|  |  | Delta | 0.4791 | 0.2297 | 0.9715 | 0.5714 | 0.139 | 0.9757 | 0.5238 | 0.1827 | 0.9983 |
|  | Mean KOOS | Percent change | 0.6429 | 0.0856 | 0.9715 | 0.5952 | 0.1195 | 0.9757 | 0.5 | 0.207 | 0.9983 |
|  |  | Delta | 0.5238 | 0.1827 | 0.9715 | 0.4048 | 0.3199 | 0.9757 | 0.4524 | 0.2604 | 0.9983 |

**Table S13: Summary of levels of inflammatory systemic biomarkers measured at baseline in serum or plasma.** Levels of sCD163, CRP, and S100A8/A9 were measured in serum, while TNFα and IL-6 were measured in plasma. N=12 patients. sCD163: soluble CD163; CRP: C-reactive protein; TNFα: tumour necrosis factor-alpha; IL-6: interleukin‑6.

| **Subject ID** | **sCD163 (ng/mL)** | **CRP (ng/mL)** | **S100A8/A9 (ng/mL)** | **TNFα (pg/mL)** | **IL-6 (pg/mL)** |
| --- | --- | --- | --- | --- | --- |
| 1 | 218.89 | 698.63 | 479.97 | 0.30 | 0.85 |
| 2 | 270.68 | 202.51 | 599.67 | 0.37 | 3.42 |
| 3 | 283.79 | 914.72 | 773.49 | 0.68 | 1.01 |
| 4 | 400.60 | 2325.87 | 984.69 | 0.40 | 0.38 |
| 5 | 327.27 | 1822.65 | 911.91 | 0.50 | 0.53 |
| 6 | 413.49 | 261.16 | 1036.80 | 0.36 | 0.21 |
| 7 | 454.91 | 181.37 | 800.07 | 0.44 | 0.32 |
| 9 | 284.05 | 137.14 | 3912.30 | 0.21 | 0.39 |
| 10 | 347.25 | 597.44 | 1984.54 | 0.24 | 1.33 |
| 11 | 300.74 | 477.76 | 2047.35 | 0.57 | 2.57 |
| 12 | 398.38 | 1498.78 | 1496.60 | 0.39 | 0.47 |
| 13 | 298.06 | 826.45 | 1181.74 | 0.25 | 1.02 |

**Table S14: Correlations between baseline levels of systemic biomarkers and changes in KOOS (percent change or delta values) at 6, 12 and 24 months.** N=12 patients. No significant correlations were observed. Spearman’s correlation. ADL: function in daily living; KOOS: knee injury and osteoarthritis outcome score. C1-2C: collagen type I and II cleavage; collagen type II cleavage; COMP: cartilage oligomeric matrix protein; CRP: C-reactive protein; CTX-II: C‐telopeptide of type II collagen; HA: hyaluronic acid; IL-6: interleukin-6; sCD163: soluble CD163; TNFα: tumour necrosis factor-alpha.

|  | | | **6 Month** | | | **12 Month** | | | **24 Month** | | |
| --- | --- | --- | --- | --- | --- | --- | --- | --- | --- | --- | --- |
| **Biomarker** | **Follow-up Score** | **Difference calculation** | **Spearman's rho** | **p value** | **FDR p value** | **Spearman's rho** | **p value** | **FDR p value** | **Spearman's rho** | **p value** | **FDR p value** |
| C1-2C | Pain | Percent change | 0.1259 | 0.6967 | 0.9828 | 0.1608 | 0.6175 | 0.9914 | 0.042 | 0.897 | 0.9820 |
|  |  | Delta | -0.0806 | 0.8035 | 0.9828 | 0.2837 | 0.3715 | 0.9914 | 0.1121 | 0.7287 | 0.9760 |
|  | ADL | Percent change | 0.0839 | 0.7954 | 0.9828 | 0.2067 | 0.5193 | 0.9914 | 0.028 | 0.9312 | 0.9820 |
|  |  | Delta | 0.0666 | 0.8372 | 0.9828 | 0.1681 | 0.6015 | 0.9914 | 0.0035 | 0.9914 | 0.9914 |
|  | Overall | Percent change | 0.2168 | 0.4986 | 0.9828 | 0.1958 | 0.5419 | 0.9914 | 0.3077 | 0.3306 | 0.9760 |
|  |  | Delta | 0.1748 | 0.5868 | 0.9828 | 0.2727 | 0.3911 | 0.9914 | 0.2394 | 0.4535 | 0.9760 |
| C2C-HUSA | Pain | Percent change | -0.007 | 0.9828 | 0.9828 | -0.042 | 0.897 | 0.9914 | -0.2098 | 0.5128 | 0.9760 |
|  |  | Delta | 0.2067 | 0.5193 | 0.9828 | 0.007 | 0.9828 | 0.9914 | -0.0841 | 0.7951 | 0.9760 |
|  | ADL | Percent change | 0.014 | 0.9656 | 0.9828 | 0.0525 | 0.8712 | 0.9914 | -0.1958 | 0.5419 | 0.9760 |
|  |  | Delta | -0.1436 | 0.6561 | 0.9828 | -0.0911 | 0.7783 | 0.9914 | -0.035 | 0.9139 | 0.9820 |
|  | Overall | Percent change | -0.0979 | 0.7621 | 0.9828 | -0.0559 | 0.8629 | 0.9914 | -0.1608 | 0.6175 | 0.9760 |
|  |  | Delta | -0.1049 | 0.7456 | 0.9828 | -0.042 | 0.897 | 0.9914 | -0.162 | 0.615 | 0.9760 |
| COMP | Pain | Percent change | -0.1119 | 0.7292 | 0.9828 | -0.3007 | 0.3423 | 0.9914 | -0.2867 | 0.3663 | 0.9760 |
|  |  | Delta | -0.2872 | 0.3654 | 0.9828 | -0.3398 | 0.2799 | 0.9914 | -0.3257 | 0.3015 | 0.9760 |
|  | ADL | Percent change | 0.1818 | 0.5717 | 0.9828 | 0.1156 | 0.7206 | 0.9914 | -0.0629 | 0.8459 | 0.9760 |
|  |  | Delta | 0.0385 | 0.9054 | 0.9828 | 0.0035 | 0.9914 | 0.9914 | -0.063 | 0.8457 | 0.9760 |
|  | Overall | Percent change | 0.049 | 0.8799 | 0.9828 | -0.0699 | 0.829 | 0.9914 | -0.1958 | 0.5419 | 0.9760 |
|  |  | Delta | -0.049 | 0.8799 | 0.9828 | -0.1189 | 0.7129 | 0.9914 | -0.2394 | 0.4535 | 0.9760 |
| CTX-II | Pain | Percent change | 0.2238 | 0.4845 | 0.9828 | 0.3986 | 0.1993 | 0.9914 | 0.1119 | 0.7292 | 0.9760 |
|  |  | Delta | 0.5079 | 0.0918 | 0.9828 | 0.5114 | 0.0893 | 0.9914 | 0.2592 | 0.4159 | 0.9760 |
|  | ADL | Percent change | 0.0699 | 0.829 | 0.9828 | 0.2102 | 0.5121 | 0.9914 | 0.014 | 0.9656 | 0.9820 |
|  |  | Delta | -0.0315 | 0.9225 | 0.9828 | 0.1506 | 0.6403 | 0.9914 | 0.1506 | 0.6403 | 0.9760 |
|  | Overall | Percent change | 0.0629 | 0.8459 | 0.9828 | 0.2098 | 0.5128 | 0.9914 | 0.1888 | 0.5567 | 0.9760 |
|  |  | Delta | 0.1329 | 0.6806 | 0.9828 | 0.3077 | 0.3306 | 0.9914 | 0.1972 | 0.539 | 0.9760 |
| HA | Pain | Percent change | 0.1259 | 0.6967 | 0.9828 | 0.0699 | 0.829 | 0.9914 | 0.0769 | 0.8122 | 0.9760 |
|  |  | Delta | 0.3573 | 0.2542 | 0.9828 | 0.063 | 0.8457 | 0.9914 | 0.1401 | 0.6641 | 0.9760 |
|  | ADL | Percent change | 0.1538 | 0.6331 | 0.9828 | 0.1331 | 0.6801 | 0.9914 | 0.021 | 0.9484 | 0.9820 |
|  |  | Delta | 0.007 | 0.9828 | 0.9828 | -0.014 | 0.9655 | 0.9914 | 0.1506 | 0.6403 | 0.9760 |
|  | Overall | Percent change | 0.007 | 0.9828 | 0.9828 | 0.0629 | 0.8459 | 0.9914 | -0.014 | 0.9656 | 0.9820 |
|  |  | Delta | 0.035 | 0.9141 | 0.9828 | 0.049 | 0.8799 | 0.9914 | 0.0141 | 0.9653 | 0.9820 |
| CRP | Pain | Percent change | 0.007 | 0.9828 | 0.9828 | -0.4196 | 0.1745 | 0.9914 | -0.2168 | 0.4986 | 0.9760 |
|  |  | Delta | -0.3608 | 0.2493 | 0.9828 | -0.4939 | 0.1027 | 0.9914 | -0.3433 | 0.2747 | 0.9760 |
|  | ADL | Percent change | 0.1399 | 0.6646 | 0.9828 | 0.0245 | 0.9397 | 0.9914 | -0.0769 | 0.8122 | 0.9760 |
|  |  | Delta | 0.0245 | 0.9397 | 0.9828 | -0.1366 | 0.6721 | 0.9914 | -0.0981 | 0.7617 | 0.9760 |
|  | Overall | Percent change | 0.0979 | 0.7621 | 0.9828 | -0.1678 | 0.6021 | 0.9914 | -0.1259 | 0.6967 | 0.9760 |
|  |  | Delta | -0.035 | 0.9141 | 0.9828 | -0.1608 | 0.6175 | 0.9914 | -0.2113 | 0.5098 | 0.9760 |
| IL6 | Pain | Percent change | 0.1329 | 0.6806 | 0.9828 | -0.042 | 0.897 | 0.9914 | 0.2238 | 0.4845 | 0.9760 |
|  |  | Delta | 0.4308 | 0.1621 | 0.9828 | 0.063 | 0.8457 | 0.9914 | 0.3292 | 0.296 | 0.9760 |
|  | ADL | Percent change | 0.1818 | 0.5717 | 0.9828 | 0.1121 | 0.7287 | 0.9914 | 0.3007 | 0.3423 | 0.9760 |
|  |  | Delta | 0.3923 | 0.2072 | 0.9828 | 0.2837 | 0.3715 | 0.9914 | 0.4518 | 0.1403 | 0.9760 |
|  | Overall | Percent change | 0.1259 | 0.6967 | 0.9828 | 0.0979 | 0.7621 | 0.9914 | 0.1748 | 0.5868 | 0.9760 |
|  |  | Delta | 0.3147 | 0.3191 | 0.9828 | 0.0629 | 0.8459 | 0.9914 | 0.2535 | 0.4266 | 0.9760 |
| S100A8/A9 | Pain | Percent change | -0.1678 | 0.6021 | 0.9828 | 0.0769 | 0.8122 | 0.9914 | 0.0699 | 0.829 | 0.9760 |
|  |  | Delta | 0.049 | 0.8797 | 0.9828 | 0.3012 | 0.3414 | 0.9914 | 0.3363 | 0.2852 | 0.9760 |
|  | ADL | Percent change | -0.1608 | 0.6175 | 0.9828 | -0.0666 | 0.8372 | 0.9914 | -0.1049 | 0.7456 | 0.9760 |
|  |  | Delta | 0.1156 | 0.7206 | 0.9828 | 0.1296 | 0.6881 | 0.9914 | 0.0666 | 0.8372 | 0.9760 |
|  | Overall | Percent change | -0.0699 | 0.829 | 0.9828 | 0.0769 | 0.8122 | 0.9914 | 0.2238 | 0.4845 | 0.9760 |
|  |  | Delta | 0.0839 | 0.7954 | 0.9828 | 0.1119 | 0.7292 | 0.9914 | 0.2465 | 0.4399 | 0.9760 |
| sCD163 | Pain | Percent change | 0.049 | 0.8799 | 0.9828 | 0.0699 | 0.829 | 0.9914 | 0.1538 | 0.6331 | 0.9760 |
|  |  | Delta | -0.2662 | 0.403 | 0.9828 | 0.021 | 0.9483 | 0.9914 | 0.1191 | 0.7124 | 0.9760 |
|  | ADL | Percent change | -0.0629 | 0.8459 | 0.9828 | -0.0666 | 0.8372 | 0.9914 | -0.1329 | 0.6806 | 0.9760 |
|  |  | Delta | -0.1576 | 0.6247 | 0.9828 | -0.2172 | 0.4978 | 0.9914 | -0.2417 | 0.4492 | 0.9760 |
|  | Overall | Percent change | 0.0699 | 0.829 | 0.9828 | 0.007 | 0.9828 | 0.9914 | 0.1399 | 0.6646 | 0.9760 |
|  |  | Delta | -0.0769 | 0.8122 | 0.9828 | 0.021 | 0.9484 | 0.9914 | 0.1197 | 0.7109 | 0.9760 |
| TNFα | Pain | Percent change | 0.6713 | *0.0168 | 0.9828 | 0.2517 | 0.4299 | 0.9914 | 0.0909 | 0.7787 | 0.9760 |
|  |  | Delta | 0.3468 | 0.2695 | 0.9828 | 0.028 | 0.9311 | 0.9914 | -0.0876 | 0.7867 | 0.9760 |
|  | ADL | Percent change | 0.3497 | 0.2652 | 0.9828 | 0.3818 | 0.2207 | 0.9914 | 0.3007 | 0.3423 | 0.9760 |
|  |  | Delta | 0.0701 | 0.8287 | 0.9828 | 0.1401 | 0.6641 | 0.9914 | 0.2102 | 0.5121 | 0.9760 |
|  | Overall | Percent change | 0.5734 | 0.0513 | 0.9828 | 0.3776 | 0.2262 | 0.9914 | 0.2168 | 0.4986 | 0.9760 |
|  |  | Delta | 0.2448 | 0.4433 | 0.9828 | 0.2098 | 0.5128 | 0.9914 | 0.1338 | 0.6784 | 0.9760 |

**Table S15: Forward and reverse primer sequences for qPCR.**

| **Gene** | **Forward** | **Reverse** |
| --- | --- | --- |
| *ACTA2* | TTTGGCTTGGCTTGTCAGGG | GGAAGCTTTAGGGTCGCTGG |
| *ANGPT1* | AGCGCCGAAGTCCAGAAAAC | TACTCTCACGACAGTTGCCAT |
| *B2M* | CTCCGTGGCCTTAGCTGTG | TTTGGAGTACGCTGGATAGCCT |
| *CCN2* | TGTGGCTTTAGGAGCAGTG | GCTACAGGCAGGTCAGTGG |
| *CCR7* | TTTTACCGCCCAGAGAGCG | AATGACAAGGAGAGCCACC |
| *CD163* | TGGACCTAATGAATTCCTCAGAAAA | ACACAGAAATTAGTTCAGCAGCA |
| *CD206* | CTACAAGGGATCGGGTTTATGGA | TTGGCATTGCCTAGTAGCGTA |
| *CD274* | TCAATGCCCCATACAACAA | TGCTTGTCCAGATGACTTCG |
| *CD86* | CTGCTCATCTATACACGGTTACC | GGAAACGTCGTACAGTTCTGTG |
| *CXCL8* | AAATTTGGGGTGGAAAGGTT | TCCTGATTTCTGCAGCTCTGT |
| *EDN1* | AGAGTGTGTCTACTTCTGCCA | CTTCCAAGTCCATACGGAACAA |
| *GAPDH* | GGAGCGAGATCCCTCCAAAAT | GGCTGTTGTCATACTTCTCATGG |
| *GUSB* | GACACGCTAGAGCATGAGGG | GGGTGAGTGTGTTGTTGATGG |
| *HLADRA* | AAGCACTGGGAGTTTGATGC | ATTGCTTTTGCGCAATCCCT |
| *HMOX1* | AAGACTGCGTTCCTGCTCAAC | AAAGCCCTACAGCAACTGTCG |
| *IL10* | CGAGATGCCTTCAGCAGAGT | CGCCTTGATGTCTGGGTCTT |
| *IL12A* | CCTCCACTGTGCTGGTTTTAT | TCAGCAACATGCTCCAGAAG |
| *IL1B* | GTACCTGTCCTGCGTGTTGA | GGGAACTGGGCAGACTCAAA |
| *PDCD1LG2* | GTACATAATAGAGCATGGCAGCA | CCACCTTTTGCAAACTGGCTGT |
| *STAB1* | GAACCATGTGCCACTGGAAGGC | AGCGGAATCTCCTGGTGCAGTT |
| *TBP* | AGCGCAAGGGTTTCTGGTTT | AATAGGCTGTGGGGTCAGTC |
| *THBS1* | AGACTCCGCATCGCAAAGG | TCACCACGTTGTTGTCAAGGG |
| *TNFAIP6* | AGCACGGTCTGGCAAATACA | ATCCATCCAGCAGCACAGAC |
| *TREM1* | AGTTGCAGCTCGGAGTTCTGAGACA | GAACCATGTGCCACTGGAAGGC |

**Table S16. Biomarkers measured in patient serum and plasma.** Serum/plasma column indicates whether the analyte was measured in serum or plasma. CRP: C-reactive protein; IL-6: interleukin-6; TNFα: tumour necrosis factor-alpha; C1-2C: C1‐2C collagen type I and II cleavage; HA: hyaluronic acid; COMP: cartilage oligomeric matrix protein; CTX-II: C‐telopeptide of type II collagen; C2C-HUSA: collagen type II cleavage; MMP: matrix metalloproteinase; TIMP1: tissue inhibitor of metalloproteinases 1; CXCL: C‐X‐C chemokine motif ligands; CCL2: Chemokine C-C motif ligand 2; VEGFA: vascular endothelial growth factor A; IL: interleukin; HGF: hepatocyte growth factor.

| **Biomarker** | **Serum/Plasma** | **Measurement method** |
| --- | --- | --- |
| S100A8/A9 | Serum | S100A8/A9 Heterodimer DuoSet ELISA (R&D Systems; Cat. DY8226-05 |
| CRP | Serum | CRP DuoSet ELISA (R&D Systems; Cat. DY1707) |
| CD163 | Serum | CD163 DuoSet ELISA (R&D Systems; Cat. DY1607). |
| IL-6 | Plasma | High-sensitivity IL-6 ELISA (ThermoFisher Scientific; Cat. BMS213HS)) |
| TNFα | Plasma | Quantikine HS TNFα ELISA (R&D Systems; HSTA00E). |
| C1-2C | Serum | Previous study [1] |
| HA | Plasma | Previous study [1] |
| COMP | Plasma | Previous study [1] |
| CTX-II | Urine | Previous study [1] |
| C2C-HUSA | Urine | Previous study [1] |
| MMP-3 | Synovial fluid | Previous study [1] |
| MMP-9 | Synovial fluid | Previous study [1] |
| MMP-13 | Synovial fluid | Previous study [1] |
| TIMP1 | Synovial fluid | Previous study [1] |
| CX3CL1 | Synovial fluid | Previous study [1] |
| CXCL1 | Synovial fluid | Previous study [1] |
| CCL2 | Synovial fluid | Previous study [1] |
| VEGFA | Synovial fluid | Previous study [1] |
| IL12p40 | Synovial fluid | Previous study [1] |
| IL-6 | Synovial fluid | Previous study [1] |
| IL-8 | Synovial fluid | Previous study [1] |
| HGF | Synovial fluid | Previous study [1] |
| sCD14 | Synovial fluid | Previous study [1] |
| sCD163 | Synovial fluid | Previous study [1] |
| prostaglandin; | Synovial fluid | Previous study [1] |
| adiponectin | Synovial fluid | Previous study [1] |
| adipsin | Synovial fluid | Previous study [1] |
| leptin | Synovial fluid | Previous study [1] |
| resistin | Synovial fluid | Previous study [1] |
