## Supplementary figures and images for "Responders vs. non-responders to mesenchymal stromal cells in knee osteoarthritis patients: mechanistic correlates of donor cell attributes and putative patient features"

### Graphical Abstract

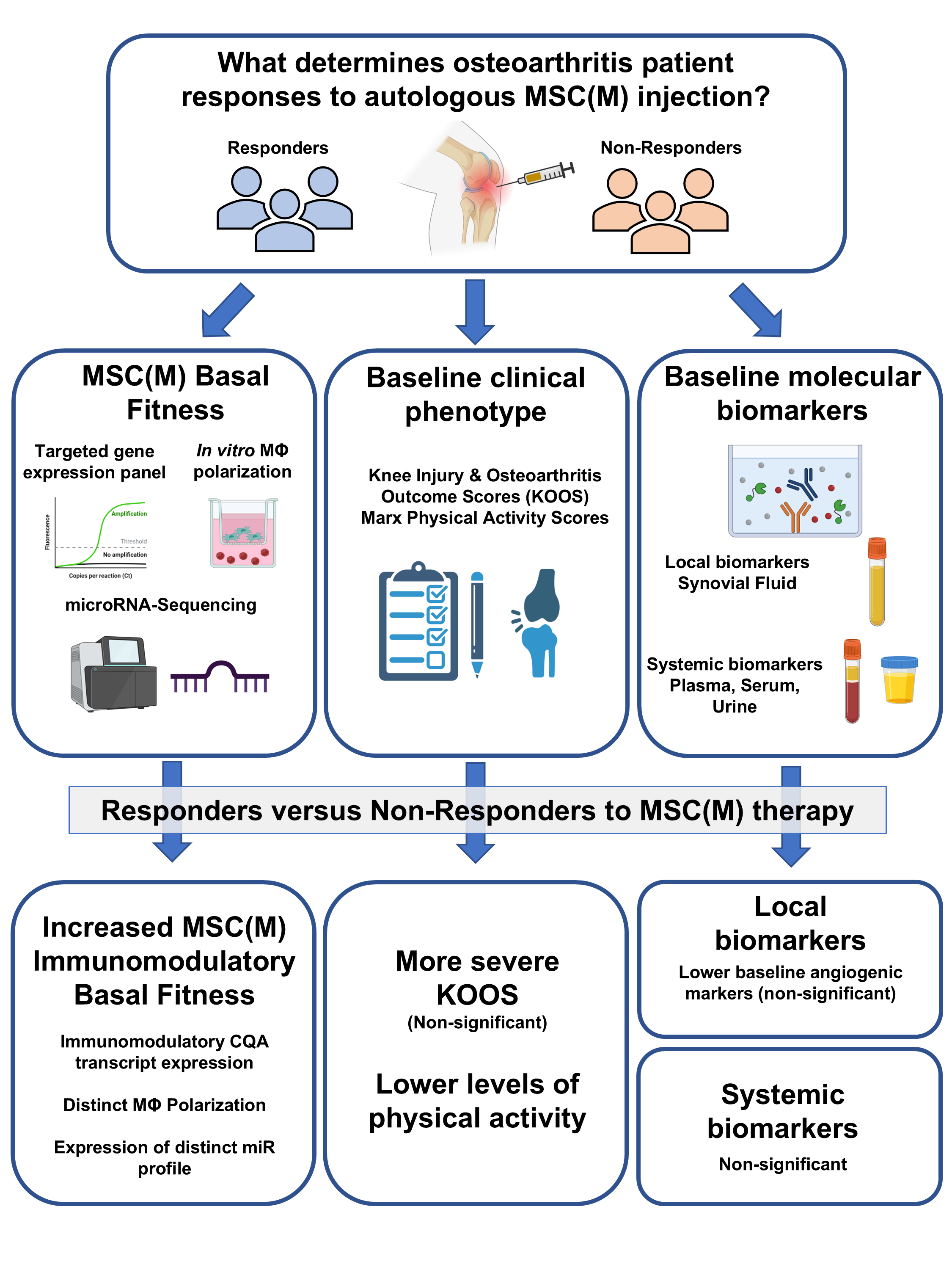
